# Thermotaxis behavior of *Drosophila melanogaster*: A quantitative analysis of sensory-motor integration and heat avoidance

**DOI:** 10.1101/2025.08.11.668339

**Authors:** Zeinab Moradi, Sara Jahedazad, Ehsan Hosseinian, Tayebeh Minaei, Ali Sadighi

## Abstract

Bridging the gap between sensory input and behavioral output remains a central challenge in neuroscience. In Drosophila, the seemingly simple behavior of heat avoidance reflects a surprisingly intricate interplay between sensory processing and rapid decision-making, one that can only be fully understood through precise stimulus control and high-resolution behavioral analysis. Here, we present an integrated platform combining a precision-controlled thermal arena with deep learning–based tracking and a suite of statistical, machine-learning, and behavioral modeling tools to quantify fine-grained locomotor dynamics across thermal gradients. Wild-type flies exhibited robust temperature-dependent strategies, including increased avoidance indices and U-turn frequencies at higher temperatures. Antennal thermoreceptor ablation disrupted rapid decision-making at thermal boundaries while preserving baseline heat avoidance, suggesting the presence of parallel thermosensory pathways. Flies with antennal impairments showed 9% higher active avoidance within hot zones, significant reductions in U-turn responses, increased traversal speeds, weakened centrophobism, and staccato movement patterns that remained unaffected. Supervised classifiers achieved 76.7% accuracy in distinguishing temperature conditions and 83.0% accuracy in identifying sensory impairments. A Braitenberg vehicle model replicated key behavioral patterns, but its reduced adaptability to abrupt changes in sensory input underscored the superior compensatory capacity of biological systems. These findings offer unprecedented quantitative resolution into Drosophila’s thermotaxis behavior and highlight the distinct contributions of peripheral and internal thermoreceptors to navigational decision-making. This scalable framework advances the study of sensorimotor integration and has potential applications in bio-inspired navigation design.

## Introduction

Navigation in animals is achieved by breaking complex tasks into smaller steps and employing sensorimotor strategies to resolve each one. Through these strategies, an animal can assess its position relative to a target location and make informed decisions to reach its destination ^1,2^. Drosophila melanogaster is one of the most genetically and physiologically tractable model organisms in biology ^3-8^. Several genes were first discovered in Drosophila and later found to have human analogs ^9^. This organism has been instrumental in studying a wide range of genes linked to human diseases, including neurodegenerative and neurodevelopmental disorders (e.g., Parkinson’s, Huntington’s, Alzheimer’s, Amyotrophic Lateral Sclerosis, Spinal Muscular Atrophy, and Autism Spectrum Disorder), infectious diseases, sleep disorders, cancer, drug addiction, cardiac failure, and diabetes ^6,7,10-16^. The broad utility of Drosophila in biomedical research extends beyond disease modeling to fundamental investigations of sensory processing and behavior. Among these, thermotactic behavior offers valuable insights into neural circuit function, sensory adaptation, and decision-making. Leveraging machine learning and behavioral modeling techniques, thermotactic responses can be analyzed in a comprehensive manner, providing a deeper understanding of the biological principles governing temperature-driven behaviors. Temperature is a critical environmental factor that directly influences the rate of biochemical reactions and biological processes, thereby affecting the physiology, morphology, and behavior of animals ^17-19^. Unlike endothermic animals, which adapt to temperature through metabolic mechanisms, ectothermic animals like Drosophila lack central temperature regulation and instead rely on behavioral strategies to stabilize their body temperature ^18,20^. The ability of ectotherms to withstand temperature fluctuations is strongly influenced by their body size. Due to its small size and large surface-area-to-volume ratio, the body temperature of Drosophila equilibrates rapidly with environmental temperatures ^20-25^. Consequently, Drosophila exhibits pronounced behavioral preferences for temperatures conducive to survival and reproduction, particularly around 25°C ^26^. To maintain their preferred temperature, they move away from excessive warmth (negative thermotaxis) or from excessive cold (positive thermotaxis) by navigating temperature gradients. Thermotaxis in Drosophila is mediated by specialized thermoreceptors that sense temperature changes and drive distinct behavioral responses. Drosophila melanogaster possesses at least three types of thermoreceptors: moderate warmth sensors, moderate cold sensors, and noxious warmth sensors, although a noxious cold sensor has yet to be identified. Thermoreception in Drosophila is achieved through three cold cells in the aristae and sacculus cells, which sense cold, and three hot cells in the aristae and anterior cells, which detect warmth ^23,27^. These thermoreceptors utilize distinct molecular mechanisms, project to different brain regions, and drive unique behavioral outcomes ^23,28-30^. For instance, Gustatory Receptors (Gr) in hot cells are essential for rapid thermotaxis in steep thermal gradients, while Transient Potential Receptor (TRP) channels in anterior cells enable heat avoidance in gradual thermal gradients ^23,31^. The differentiation in thermoreceptor functionality provides Drosophila with a robust mechanism for adapting to varying thermal environments. The thermal response of Drosophila is further modulated by social dynamics, with group behavior playing a significant role in navigating harmful thermal stimuli. Individual behavior in Drosophila is often influenced by group size, local density, and interactions within the group ^32^. Rooke et al. ^32^ emphasized that Drosophila can perceive the number of individuals in its group, with social interactions enhancing environmental sensing and decision-making efficiency ^33^. These dynamics influence behaviors such as foraging ^2^, circadian rhythms ^34^, and odor avoidance ^33^. Group locomotion often involves cascades of appendage-touch interactions, which enhance collective thermal avoidance behavior. Analyzing group locomotion behavior in thermal preference assays provides valuable insights into the interplay between individual and collective decision-making processes. To investigate thermotactic behavior in Drosophila, researchers have employed a variety of behavioral assays. Devices for studying thermotaxis can be broadly categorized into temperature gradient-based setups and two-choice assays ^35^. In gradient-based assays, animals are placed on a plate with a temperature gradient, and their positional preferences are analyzed based on metrics such as time spent in specific temperature zones and distribution across the gradient ^36^. Linear temperature gradients are most commonly employed, with methods ranging from the use of aluminum slabs with hot and cold water at opposite ends ^37^ to thermoelectric coolers and resistive heaters for precise temperature control ^1,38^. One significant disadvantage of gradient-based assays is that they may not fully capture rapid or binary thermotactic decision-making in animals. While gradient assays provide valuable information about long-term thermal preferences and distribution across a range of temperatures, they are less suited for studying immediate, discrete choices between two distinct temperatures. This limitation arises because the gradient creates a continuous spectrum of temperatures, which may not reflect real-world scenarios where animals often encounter and must choose between two distinct thermal environments. Additionally, the interpretation of positional preferences in gradient assays can be complex, as factors such as exploration behavior, acclimatization time, and gradient steepness can influence the results. To address these limitations, two-choice assays offer a complementary approach by presenting animals with a clear binary decision between two distinct temperatures. This method is particularly useful for studying rapid thermotactic responses and the underlying neural mechanisms of temperature preference, as it simplifies the decision-making process and allows for more straightforward interpretation of behavioral outcomes. A critical aspect of such experiments is the precise control of temperature. Researchers may employ various methods, including thermoelectric modules, melting substances, or digital temperature controllers like PID systems, depending on the desired precision. Building on the advantages of precise temperature control demonstrated in previous studies ^39^, in this work a versatile temperature preference device is designed utilizing Peltier modules, controlled via a microcontroller board and implemented a PID algorithm for precise thermal regulation. This approach ensures high accuracy and stability in maintaining desired temperatures, which is critical for reliable thermotactic experiments. In addition to designing assays, accurately quantifying animal behavior is essential for interpreting thermotactic responses. Visual tracking of small organisms like Drosophila provides a cost-effective and noninvasive method for analyzing behavioral data with high spatiotemporal resolution ^40-42^. Advances in computer vision have facilitated the development of automated tracking tools, enabling reproducible, high-throughput behavioral analyses ^33,43-52^. These tools enhance replicability and minimize observer bias ^51^, allowing for a wide range of detailed analyses of behaviors such as locomotion ^43-45,53-55^, social interactions ^2,32-34,46,54,56-58^, and gene-neuron-behavior relationships ^4- 7,10,13,29,30,39,44,48,54,56,59-63^. By integrating behavioral assays with automated tracking, this study aims to provide a comprehensive understanding of thermotactic behavior in Drosophila melanogaster. This work not only highlights the intricate mechanisms underlying thermo-sensation but also underscores the value of Drosophila as a model organism for investigating complex biological phenomena.

In this study, we have developed a versatile experimental framework to investigate the negative thermotactic behavior of Drosophila melanogaster, leveraging advanced technologies and computational tools to deepen our understanding of thermal navigation. By designing a two-choice temperature preference assay with Peltier controlled modules, real time stable thermal regulation is achieved, enabling the study of rapid thermotactic responses under controlled conditions. Complementing this setup, we implemented a state-of-the-art video-tracking workflow using YOLOv5 and DeepSORT algorithms, which allowed for the extraction of detailed behavioral parameters such as speed, avoidance index, and centrophobism. These tools provided valuable insights into the role of temperature receptor neurons (TRNs) in the antennae and their contribution to heat avoidance strategies. Additionally, fly behavior was simulated using a Braitenberg vehicle model, demonstrating the simplicity and elegance of sensory-motor transformations underlying thermotaxis. These findings align with and extend previous literature while highlighting the complexity of thermal navigation and emphasizing the interplay between peripheral and internal thermosensory mechanisms.

## Results

### Spatial Maps of Temperature Preference

Navigation of the desired area manifests the fruit fly’s temperature preferences. Thus, the degree of desire to be present in a region can be a useful indicator of avoidance intensity. To visualize the temperature preferences in the circular chamber, we have drawn location intensity (percentage of presence) maps. Since long stops (speeds below 1 mm/s) aren’t taken into account on these maps, they are representing the active percentage of presence.

Location intensity maps for the experiments with base temperatures in quadrants I & III are shown in *Figure 1 – a*. It is evident from the maps that at 25°C, the distribution of flies in the circular chamber is less different from uniformity, but with increasing temperature stimulation, this difference increases and heat avoidance behavior intensifies. This observation is further supported by the results of the Kolmogorov-Smirnov test applied to the angular position data, where the test statistic increased from 0.06 at 25°C to 0.41 at 40°C, with a 95% confidence interval (*Figure 1* – a). Also, it seems that at 40°C, a virtual boundary surrounds the flies in the quadrants with the base temperature, preventing them from entering the quadrants with the test temperature. In addition, the tendency to walk along the arena’s wall is apparent in all the temperature preference assays, confirming a distinctive feature of fly’s locomotive behavior termed “centrophobism: center avoidance due to fear or anxiety” or “thigmotaxis: the attraction to the touch of the arena wall” in the corresponding literatures ^4,43,44,64^. These precious observations are considered as a starting point for more precise investigations of the locomotor behavior of fruit flies under the influence of thermal stimulation. The percentage of presence maps at test temperature of 25°C and 35°C for antennae-ablated group of flies is displayed in *Figure 1 – a*. However, in comparisons between control and manipulated groups at 35°C, antennae-ablated type exhibits an increase in active presence in warm quadrants, which can be interpreted as a decrease in their heat-avoidance behavior. As confirmed by the Kolmogorov-Smirnov test (*p*<0.001), wild and antennae-ablated groups have marked differences in angular position distribution. The results remain consistent for experiments conducted at base temperatures within the quadrants II & IV, accessible in the supporting materials.

**Figure 1.**
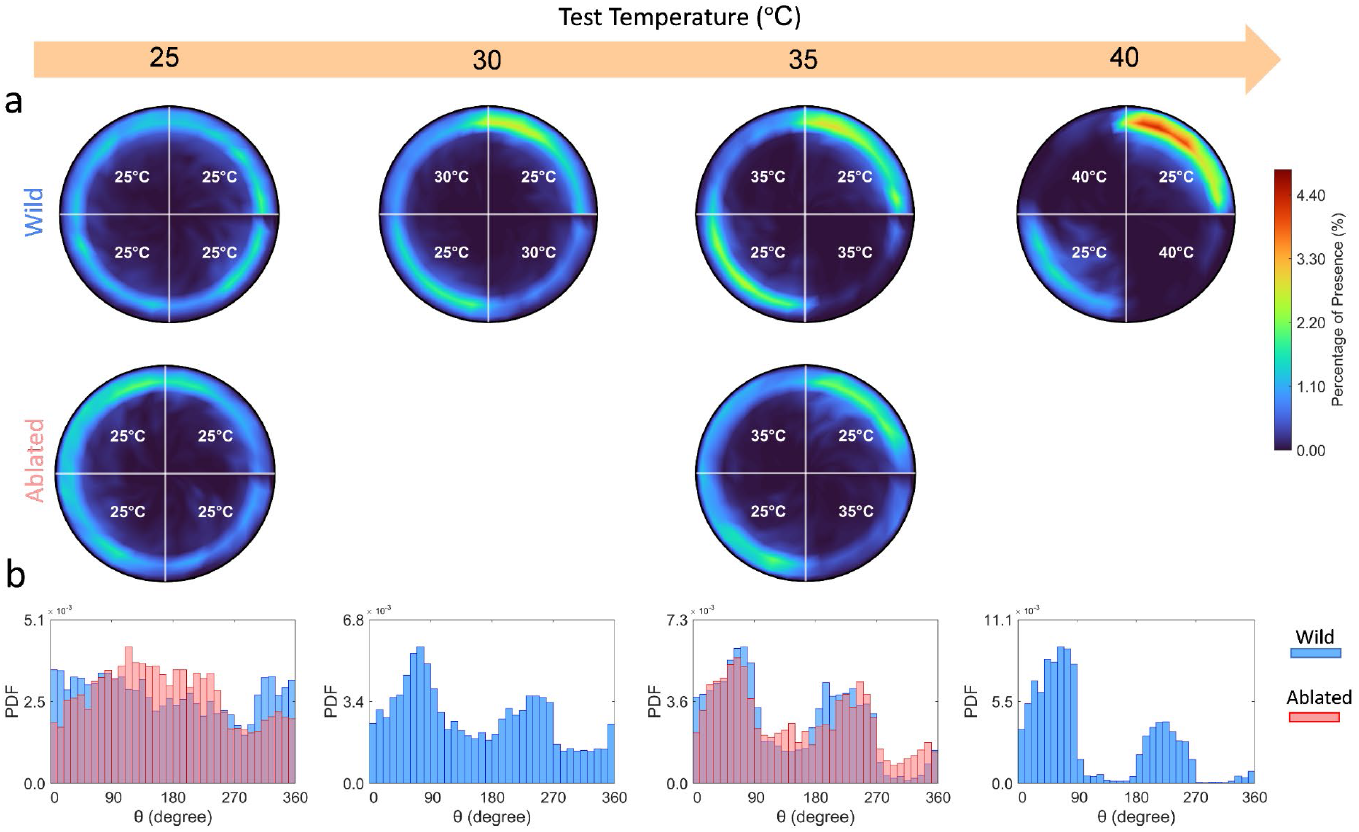
Spatial Maps of Temperature Preference. (a) Percentage presence maps of wild-type and antennae-ablated groups during two-choice temperature preference assays for test temperatures of 25°C, 30°C, 35°C, and 40°C on quadrants II and IV. These maps reveal Drosophila’s collective temperature preference. The results of the experiments with the reversed spatial configuration of the base and test quadrants are provided in the supporting materials. Wall-following behavior of flies, known as centrophobism, is apparent under all thermal conditions. Heat avoidance increases with temperature, especially at 40°C, where flies avoid test quadrants entirely, suggesting a virtual boundary. Antennae-ablated flies show higher activity in warmer zones at 35°C, indicating reduced thermal aversion. (b) Probability density function of angular position confirms greater heat avoidance with increasing test temperatures. The tendency to choose the secondary diagonal quadrants (θ=[0°,90°] and θ=[180°,270°]) with a desired temperature of 25°C is strengthened when the test temperature is raised from 25°C to 40°C. The variability in the distribution of flies observed in the favorable quadrants I and III can likely be attributed to inaccuracies in temperature regulation within the experimental device, as well as the intricate dynamics of collective behavior within the groups. Prior research ^32^ indicates that individual flies have an awareness of the number of others in their group, as both the overall group size and local nearest-neighbor density alter their behavior.

**Figure 2.**
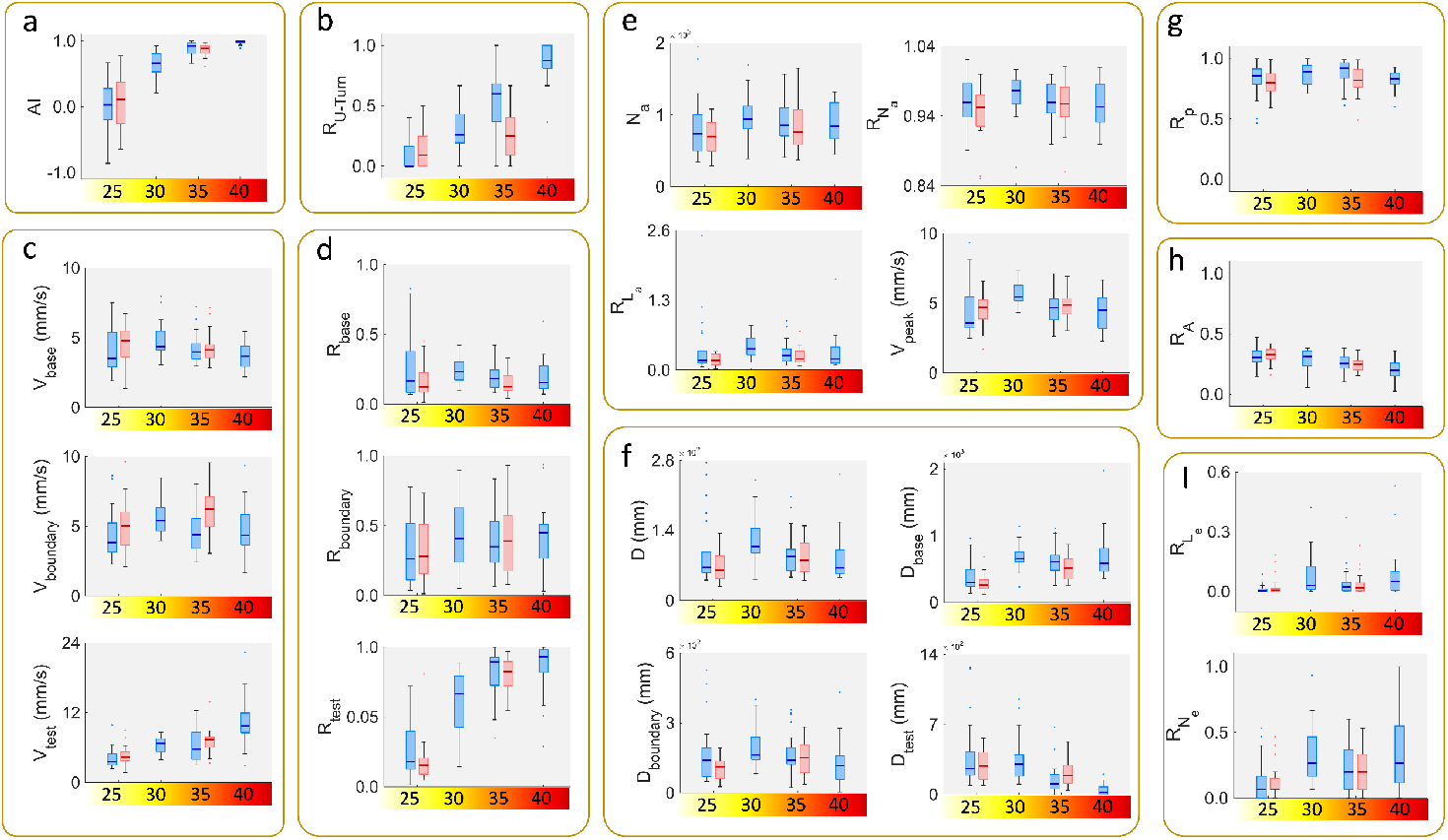
Statistical Analysis of Locomotor Parameters. For the wild-type groups, shown with blue boxes, the number of assay replicates at each test temperature was as follows: n_25_=21, n_30_=20, n_35_=28, n_40_=21. Depending on the data distribution and variance assumptions, group means were compared using appropriate statistical hypothesis tests, including One-way ANOVA, Welch ANOVA, and the Kruskal–Wallis test. The resulting *p*-values for key locomotor parameters are summarized below: (a) Avoidance Index (AI): *p*<0.001; (b) Ratio of U-turns (R_U-turn_): *p*<0.001; (c) Average speed of activity in each region (V_base_, V_boundary_, V_test_): *p*_base_=0.026, *p*_boundary_=0.11, *p*_test_<0.001; (d) Time ratio of activity in each region (R_base_, R_boundary_, R_test_): *p*_base_=0.22, *p*_boundary_=0.27, *p*_test_<0.001; (e) the *p*-values for the number of active episodes and its frequency ratio and length ratio 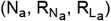 are 0.71, 0.51, 0.50, respectively and for the peak speed (V_peak_) is 0.0012; (f) Total distance traveled (D): *p*=0.173; Distance traveled within the base, boundary, and test regions (D_base_, D_boundary_, D_test_): *p*_base_=0.0028, *p*_boundary_=0.18, *p*_test_<0.001; (g) Time ratio spent in the peripheral area (R_p_): *p*=0.092; (h) Ratio of covered area (R_A_): *p*<0.001; (I) Number index (frequency) of pairwise encounters 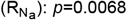 and time index of pairwise 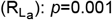. For the manipulated (antennae-ablated) groups, the number of replicates was: n_25_=21, n_35_=25. To assess the role of antennal receptors in thermally modulated locomotion, locomotor parameters were compared between the wild-type and antennae-ablated groups at 35°C using either an independent t-test (assuming equal or unequal variance) or the Mann–Whitney U test. The results, indicated by red boxes, are as follows: (a) AI: *p*=0.23; (b) R_U-turn_: *p*<0.001; (c) V_base_, V_boundary_, V_test_: *p*_base_=0.69, *p*_boundary_<0.001, *p*_test_=0.32; (d) R_base_, R_boundary_, R_test_: *p*_base_=0.096, *p*_boundary_=0.41, *p*_test_=0.82; (e) 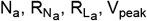: *p*-values are 0.6, 0.62, 0.29, and 0.75 respectively; (f) D: *p*=0.049; D_base_, D_boundary_, D_test_: *p*_base_=0.036, *p*_boundary_=0.44, *p*_test_=0.13; (g)) R_p_: *p*=0.0285; (h) R_A_: *p*=0.46; (I) 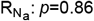 and 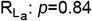.

### Statistical Analyses

The statistical hypothesis tests, as outlined in the methodology section, were employed to assess whether the means of the groups differed significantly. A highly significant increase in AI was observed in the wild-type group when the temperature was raised from 25°C to 40°C (Welch ANOVA: *p*<0.001). Similarly, the ablated-type group showed comparable results between 25°C and 35°C (t-test with unequal variance: *p*<0.001). Interestingly, when comparing the wild-type and ablated-type groups at 35°C, no significant difference in AI was detected, with the wild-type group exhibiting slightly higher AI values (t-test with equal variances: *p*=0.23). This minor difference may be explained by the presence of other thermoreceptors, in addition to the hot-activated receptors of the antenna (referred to as “Hot Cells” or HCs) ^23,28-30,39^, such as internal heat sensors in the head capsule (“Anterior Cells” or ACs) and multi-modal thermal/mechanical nociceptors in the fly’s body, which contribute significantly to heat avoidance behavior. Regarding the number of active and inactive episodes, as well as their count ratio (frequency) and length ratio (temporal ratio), no statistically significant differences were found between groups, regardless of the test temperature or antennae status. These results suggest that the overall movement pattern of the fruit fly group remains consistent, characterized by short bursts of fast movement followed by long periods of slower movement, irrespective of the test temperature or antennae manipulation. The results also show notable changes in the average peak speed of activity episodes in the wild-type group (Welch ANOVA: *p*=0.0012), but a slight statistical difference was observed when comparing the wild and ablated groups at 35°C (t-test with equal variances: *p*=0.75). The “Ratio of U-turns” was identified as a crucial metric for understanding temperature boundary decision-making strategies. Upon encountering the cool or hot boundary in the arena, control flies typically perform sharp U-turns and immediately return to the 25°C quadrant. This behavior is most prevalent at the 25°C/40°C boundary, where control flies seem almost entirely confined to the 25°C quadrant, as if restricted by an invisible barrier (Welch ANOVA: *p*<0.001). In contrast, antenna-ablated flies showed a significant reduction in this U-turn ratio at 35°C compared to the control group (t-test with equal variances: *p*<0.001), indicating that these flies adopt different strategies to avoid noxious heat. Specifically, antenna-ablated flies were less likely to perform U-turns and more frequently entered the hot quadrants. The temporal ratio spent in the peripheral area (from 0.65R_C_to R_C_) was assessed as a measure of centrophobic behavior. Statistical analysis revealed that at 35°C, the presence or absence of antennae significantly influenced the time spent in the peripheral area, with antenna-ablated flies spending less time in the periphery compared to the wild-type group (t-test with unequal variances: *p*=0.029). The mean and median differences between the groups were approximately 7% and 10%, respectively. The “Temporal Ratio of Activity,” representing the proportion of time spent in an active state (i.e., moving at speeds above 1 mm/s), demonstrated that in the wild-type group, the active presence in the test region increased with temperature. Moreover, for all test temperatures above 25°C, the active ratio was greater in the test region than in the boundary region, and the boundary region had a higher active ratio than the base region. Analysis of the average speed of movement in the base and test regions revealed that temperature significantly affected the movement speed of the wild-type group (Welch ANOVA: *p*_base_=0.026, *p*_boundary_=0.11, *p*_test_<0.001). In contrast, the behavior of antenna-ablated flies was disturbed, as they were more likely to cross into the thermal boundary (indicating a lower U-turn ratio) but exhibited higher speed when traversing from the base quadrant to the hot quadrant. The average speed in the boundary region for the manipulated flies was significantly higher than that of the wild-type group, increasing from 4.63 mm/s to 6.17 mm/s (t-test with equal variances: *p*<0.001).

Changes in the distance traveled in the test, base, and boundary regions were also observed, with increasing temperature leading to a greater preference for the base area and a decreased desire to remain in the test area. This was confirmed by the Welch ANOVA test for the wild-type group, which yielded *p*_base_=0.0028, *p*_test_<0.001. When comparing the wild-type and ablated groups at 35°C, a significant difference was found in the base region, with the wild-type group covering a greater distance (t-test with equal variances: *p*=0.036).

Lastly, measuring the area covered by the flies provided an estimate of their dispersion within the chamber. As temperature increased, the wild-type group displayed a noticeable decrease in area covered (1-way ANOVA: *p*<0.001), as the flies tended to avoid the temperature boundaries and concentrate in the base area. However, the difference between the wild-type and manipulated groups was not statistically significant (t-test with equal variances: *p*=0.46). Both the frequency and duration of pairwise encounters increased with rising temperature for both groups.

### Classification of Thermal Conditions and Deficits in Antennal Thermosensory Inputs

To gain a clearer understanding of the contribution of each parameter in distinguishing thermal conditions and impairments in antennal thermosensory inputs, we conducted a feature ranking analysis using the p-values in 1-way analysis of variance test, where the scores in the *Figure 3 – c* correspond to -log(*p*). Applying dimension reduction by principal component analysis (PCA) allows a better visualization of the data for temperature and antennae-status classification. The first three components explain 71.50% and 66.58% of all variability, respectively. Furthermore, 99% of the variability is covered by 16 and 15 components, respectively. Among all the trained models in predicting four temperature classes, the Bag ensemble classifier achieved the best performance with 76.7% accuracy. The properties of this model are as follows: number of learners=439, maximum number of splits=11, and number of predictors to sample=11. This classifier can distinguish 25°C, 30°C, 35°C, and 40°C temperature classes with sensitivity (TPR) of 90.5%, 70%, 67.90%, and 81.00%, respectively. The values of the area under the curve (AUC) of this classifier for four temperature classes are equal to 0.974, 0.834, 0.837 and 0.959 respectively, which indicates the proper performance in separating the temperature classes. However, if the volume of the training data set is increased, it is predicted that the classification accuracy will also improve. The exact process was done for antennae status discrimination. The best performance was obtained by the support vector classifier with 83% accuracy. The details of this model are as follows: kernel function of gaussian, multiclass method of one-vs-one, and box constraint level of 6.0037. The AUC value for this classifier is 0.8096.

**Figure 3.**
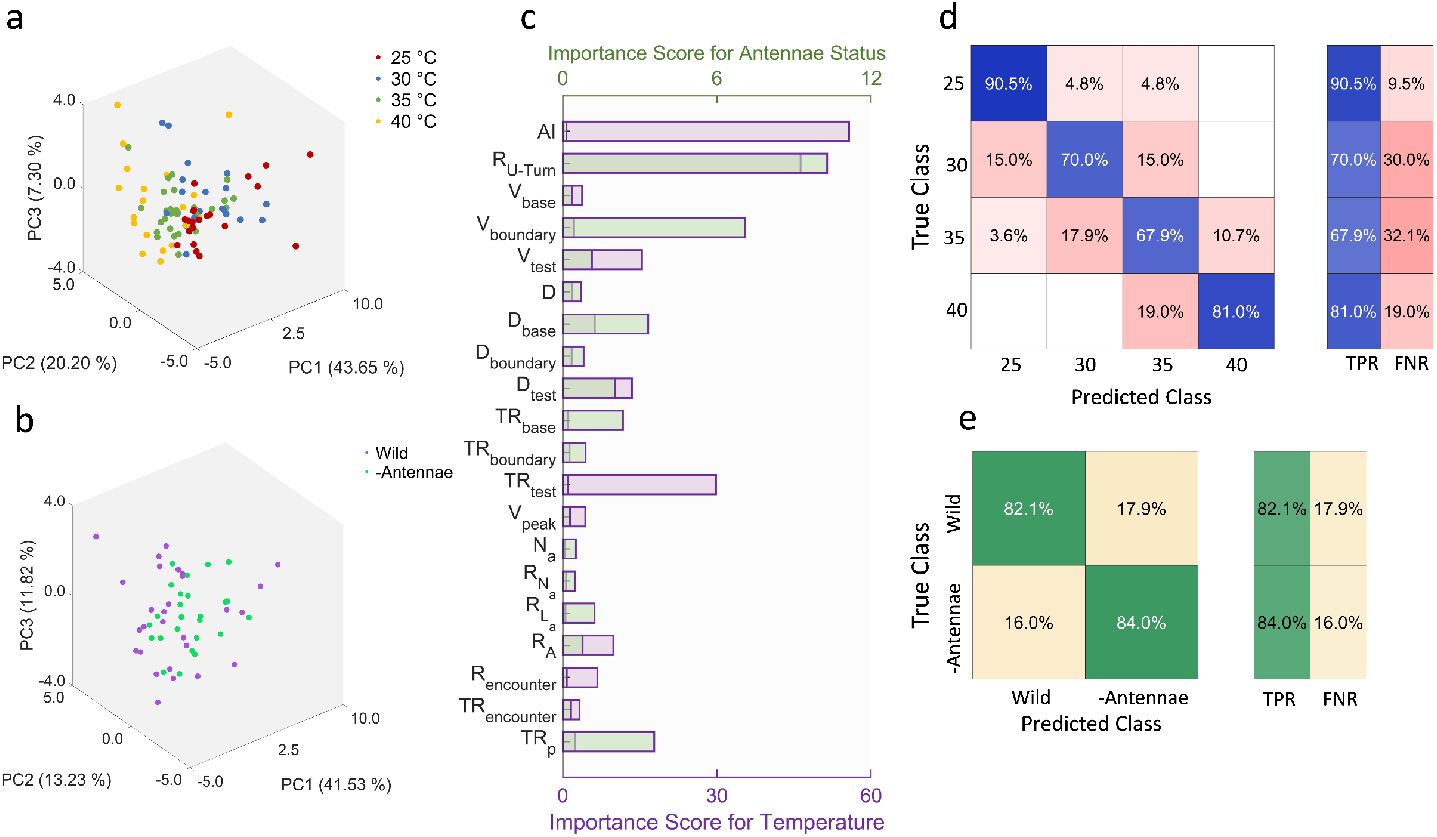
Classification on Thermal Conditions and Impairments in Antennal Thermosensory Inputs. (a, b) Principal component analysis (PCA) applied to datasets grouped by temperature (a) and antennae status (b). The first three components captured 71.5% and 66.6% of the total variance, respectively. (c) Feature ranking based on *p*-values from a one-way ANOVA; scores are represented as -log(*p*), highlighting the key parameters that contribute most significantly to differences between groups. (d, e) Confusion matrices showing classification performance. (d) Temperature classification of the control group across four conditions (25°C, 30°C, 35°C, 40°C), with the Bagg ensemble classifier achieving the highest accuracy (76.7%).(e) Classification between wild-type and antennae-ablated flies at 35°C, where the Support Vector Machine (SVM) reached 83% accuracy.

### Simulation of Thermotaxis Behavior Using a Braitenberg Vehicle

The findings suggest that the fly calculates an effective turn trajectory away from heat, utilizing sensory data from its antennae—a behavior that mirrors the “Braitenberg Vehicle” concept, a foundational model in sensorimotor transformation. Introduced by Valentino Braitenberg in 1984 ^65^, this model serves as a metaphor for understanding how complex cognitive processes can emerge from simple designs in biological systems ^66^. The Braitenberg vehicle model, as explained in the methodology section, simulates the fly’s thermotactic behavior in a 2-choice temperature preference assay at 35°C/25°C. In this simulation, a vehicle with two temperature sensors is compared to a sensor-ablated version, in which temperature input is removed entirely.

The methodology outlines the process of determining the weight coefficients (w_I_ and w_C_) based on the region, with the gain (a) and offset (b) as constants independent of location. Specifically, the model identifies the vehicle’s position within the circular chamber at any given time, applying region-specific coefficients to convert the sensed temperature into wheel speed. Additionally, the initial speed in each region (test, base, or boundary) is adjusted based on experimental observations. The parameter values are provided in the supporting materials. The results demonstrate that this model effectively generates heat-avoidance behaviors similar to those observed in real flies.

While acknowledging that the experimental results stem from group assays where inter-fly interactions play a significant role in thermotaxis responses, the goal of this model is not to replicate every aspect of the experimental scenario. Instead, it highlights that simulating complex behaviors through simple sensory-motor transformations is plausible, and the data provides support for this claim.

*Figure 4 – b* presents four sample trajectories generated by the in-silico vehicle during 1.5 minutes in the simulated arena, where the fly chooses between 25°C and 35°C. The tracks clearly illustrate the model’s ability to replicate the fly’s heat-avoidance behavior, with the vehicle avoiding the test region by performing sharp U-turns at the thermal boundary. To visualize the overall behavior, the percentage of presence in the circular chamber for all 20 simulated trajectories is shown. The vehicle predominantly avoids the test region, spending most of its time in the base area, particularly quadrants II & IV.

**Figure 4.**
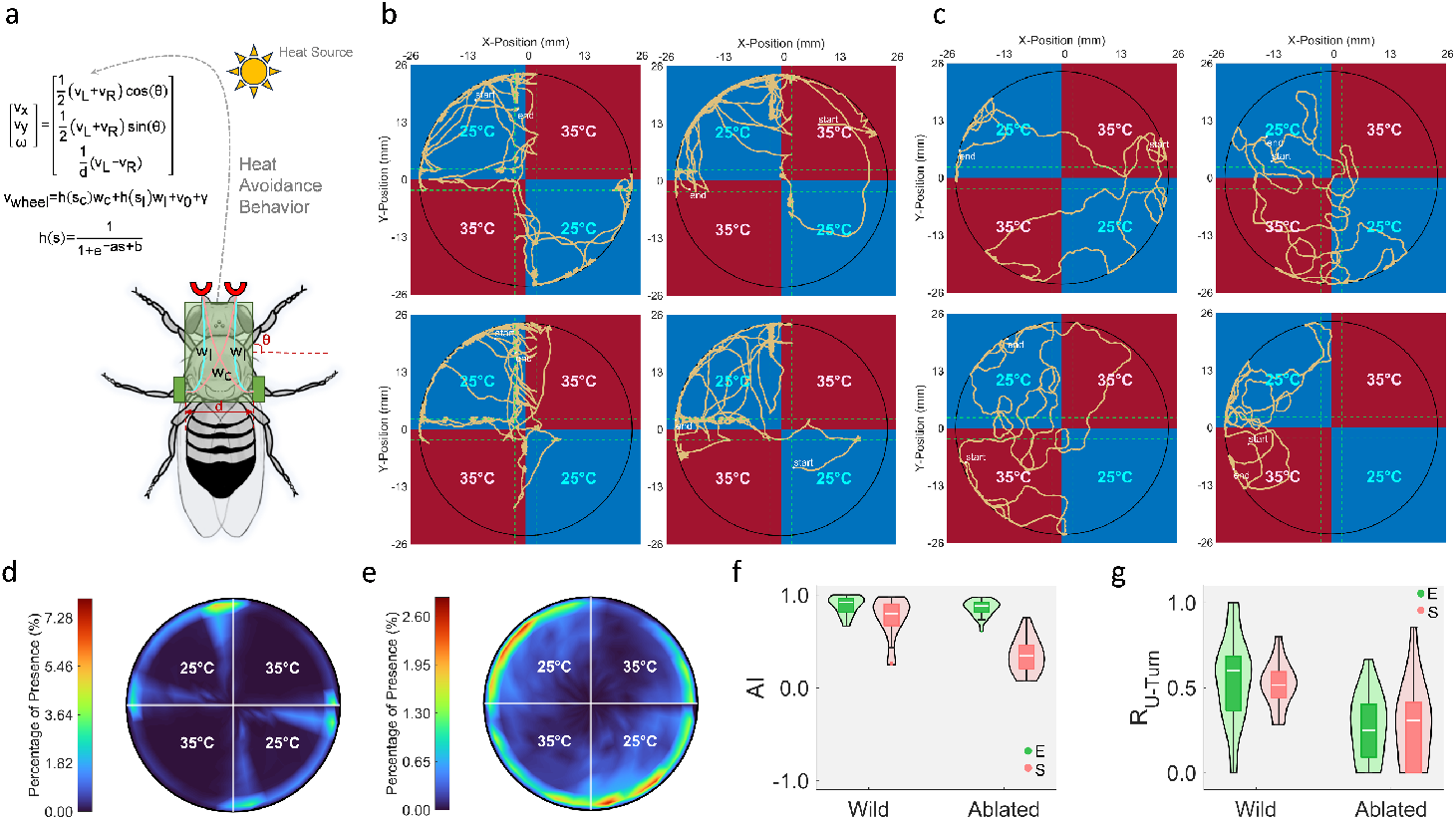
Braitenberg Vehicle Model Replicates Wild-Type Fly Thermotaxis but Exhibits Reduced Adaptability to Sensory Changes from Antennal Ablation. (a) The Braitenberg vehicle model ^65^ demonstrates how complex behaviors can arise from simple symmetric sensor–motor wiring. (b, c) Sample 1.5-minute trajectories in the simulated arena navigating between 25°C and 35°C illustrate replication of wild-type fly thermotaxis. (d, e) Location intensity maps depict the percentage of time spent in different chamber regions across simulations. (f) Avoidance Index (AI) and (g) Ratio of U-turns at 35°C compare simulated (S) and experimental (E) data; Comparisons between a wild-type fly and an in-silico vehicle show substantial consistency (*p*_AI_=0.06, *p*_RU-turn_=0.25). However, a significant difference in AI is observed when comparing antennae-ablated flies and sensor-ablated vehicles, indicating the model’s heightened sensitivity to thermal input (*p*_AI_<0.001, *p*_RU-turn_=0.97). Probability density estimates were calculated using a normal kernel function at equally spaced points across the data range.

**Figure 5.**
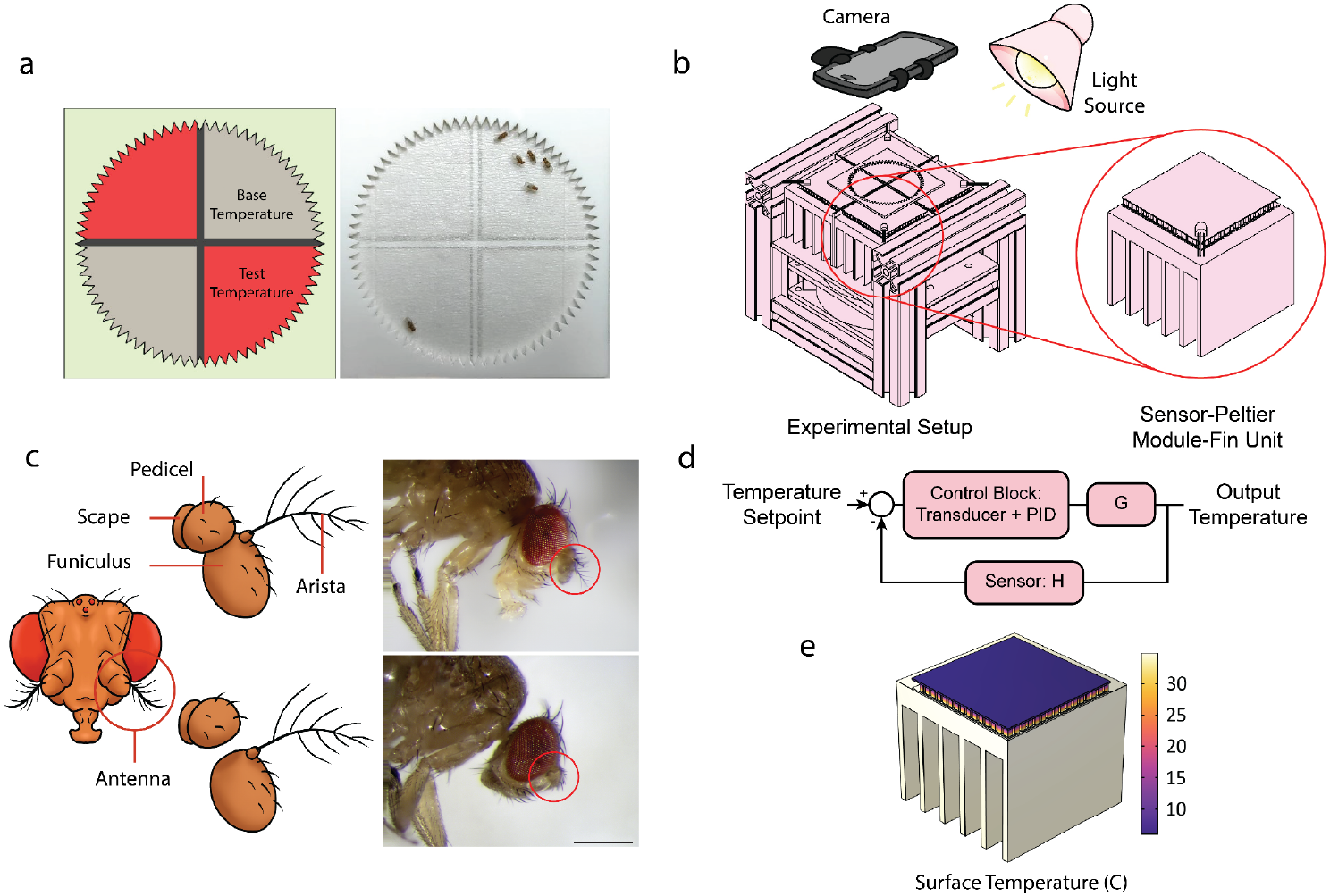
Experimental Setup and Control of the Two-Choice Temperature Preference Assay. (a) Schematic and real views of the two-choice temperature preference assay used to evaluate thermotaxis behavior in flies. (b) CAD rendering of the complete setup showing four Peltier module-sensor-heatsink units arranged in a 2×2 layout to generate distinct thermal zones. A magnified view highlights key thermal control components. The glass ceiling and wiring are omitted for clarity. (c) Antennae ablation procedure: the arista and funiculus are removed. The top row shows a fly with intact antennae; the bottom row includes a microscope image of an antennae-ablated fly. Scale bar = 500 μm. (d) Block diagram of the closed-loop control system showing the the system identification process for G, which represents the combined dynamics of the actuator (Peltier module) and other plant elements, such as fins. (e) Finite element analysis showing surface temperature distribution of a Peltier module-sensor-heatsink unit in cooling mode.

To assess the similarity between experimental and simulated results for the 25°C/35°C temperature preference assay, two metrics were used: the avoidance index (AI) for the 35°C test temperature and the ratio of U-turns/crosses at the 25°C/35°C boundary. The Kolmogorov-Smirnov test, with a 95% confidence level, confirms that both metrics share the same distribution between the experimental and simulated results (*p*_AI_=0.06, *p*_RU-turn_=0.25).

In contrast, the sensor-ablated vehicle was tested by setting the initial sensory inputs (temperature at left and right sensors) to zero (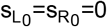 in *Figure 4 – a*). The generated trajectories of this manipulated vehicle, shown in *Figure 4 – c*, reveal that the decision-making process is disrupted, with the vehicle crossing the thermal boundary and sometimes entering the test region. The Kolmogorov-Smirnov test indicates a significant difference in the distribution of AI between the sensor-ablated vehicle and the antenna-ablated fly (*p*<0.001), although their U-turn ratios are statistically similar (*p*=0.97). The observed difference in AI between the simulated and experimental results can be attributed to the presence of internal heat sensors in the fly’s anterior cells (AC) and nociceptors in the fly’s head and body, which contribute significantly to thermotactic responses in the presence of harmful temperatures.

Although the antennae of the fly were removed in the experiments, other internal sensory mechanisms allow the fly to maintain thermotactic responses. In contrast, the Braitenberg vehicle model assumes that a sensor-ablated fly is equivalent to a vehicle with zero input from its temperature sensors, which accounts for the discrepancy in thermotaxis behavior observed in the AI. This comparison highlights the superior adaptability of the fly to changes in its sensory input state following antennae removal, compared to the in-silico vehicle.

## Discussion and Conclusions

The cumulative probability density estimate of angular position in the warm quadrants showed a significant decrease from 0.5 at 25°C to 0.06 at 40°C, indicating an intensification of heat avoidance behavior with rising temperature. This effect creates a virtual boundary at 40°C that prevents flies from entering the warmer quadrants. The cumulative probability density estimate of radial position outside the central zone was 0.86 ± 0.04, further supporting the prevalence of wall-following behavior across all thermal conditions. To rigorously assess the variation in behavioral parameters in response to temperature, statistical hypothesis tests, including One-Way ANOVA, Welch ANOVA, and Kruskal-Wallis tests, were employed. Both the “Avoidance Index” and the “Ratio of U-turns” exhibited significant increases with higher test temperatures, consistent with findings from previous influential studies. Additionally, the behavior of a manipulated group of flies with ablated antennae was examined during two-choice preference assays at 25°C and 35°C. The percentage of presence maps revealed that at 35°C, antennae-ablated flies exhibited 9% greater active presence in the warm quadrants, suggesting a diminished ability to avoid noxious heat. Despite this, the Avoidance Index remained statistically unchanged, indicating that while antennae-ablated flies can still detect heat through their anterior cells (ACs) by staying more in base regions, they rely on their antennae for rapid thermotactic responses at temperature boundaries. This reliance was further supported by a noticeable reduction in U-turn actions among the antennae-ablated flies. Interestingly, although antennae-ablated flies were more likely to cross thermal boundaries, they displayed higher traversal speeds and exhibited a reduction in centrophobism, spending more time in the central zone compared to the wild-type flies. The characteristic staccato movement pattern of Drosophila, marked by short bursts of rapid motion interspersed with prolonged slow-speed movements or complete stops, was preserved regardless of temperature conditions or antennae status.

The comprehensive dataset enabled the classification of the control group based on test temperature (25°C, 30°C, 35°C, 40°C) with 76.7% accuracy, as well as the classification of antennae status (wild-type or ablated) at 35°C with 83% accuracy. This supports the claim that the heat-avoidance behavior in fruit flies during the two-choice temperature preference assays was effectively described. The top three crucial characteristics for temperature classification included the “Avoidance Index,” the “Ratio of U-turns,” and the “Time Ratio of Activity in the Test Region.” For distinguishing fly groups based on antennae presence, key traits were the “Ratio of U-turns,” “Average Speed in the Boundary Region,” and “Time Ratio Spent in the Peripheral Area.”

In the context of the Braitenberg vehicle model, which included two temperature sensors and two motor-driven wheels, the in-silico vehicle demonstrated behavior closely resembling that of wild flies in terms of the “Avoidance Index” and “Ratio of U-turns” during the 25°C/35°C two-choice temperature preference assay. This finding suggests that a relatively simple set of rules could govern responses to a heat stimulus, with Braitenberg’s vehicle serving as one of the simplest theoretical models for sensory-motor transformation. The heat response of a sensor-ablated vehicle revealed heightened sensitivity to sudden input changes, where the distribution of the “Avoidance Index” significantly differed from that of a real antennae-ablated fly. This discrepancy highlights additional layers of complexity, which can be attributed to the presence of internal heat sensors within the head capsule (ACs), as well as thermal and mechanical nociceptors in the fly’s body, which contribute to its response to harmful temperatures. Moreover, the fruit fly’s enhanced cognitive capacity for learning and memory surpasses the capabilities of the Braitenberg vehicle model, further emphasizing the complexity of real-world thermotactic behavior.

This study reveals several fundamental aspects of Drosophila thermotactic behavior through rigorous quantitative analysis. The data demonstrate that centrophobic behavior (wall-following tendency) shows significant reduction in antennae-ablated flies (*p*=0.0285), with mean peripheral activity decreasing by 7-10%, establishing a novel connection between antennal input and spatial navigation strategies that extends beyond simple heat avoidance. Analysis of locomotor patterns reveals the remarkable stability of the staccato movement rhythm across all experimental conditions. The characteristic alternation between active bursts (>1 mm/s) and inactive periods persists consistently regardless of temperature (25-40°C; Welch ANOVA *p*>0.05 for all comparisons) or antennal status (t-test *p*=0.75 for peak speed differences), suggesting this fundamental pattern emerges from hardwired circuits independent of thermosensory modulation.

Crucially, the study uncovers a dissociation in thermosensory function through detailed kinematic analysis. While overall avoidance indices remain comparable between intact (AI=0.62±0.11) and antennae-ablated flies (AI=0.59±0.13; *p*=0.23) at 35°C, examination of active movement periods reveals ablated flies spend 9% more time in warm zones (*p*<0.01) and show 33% higher boundary crossing speeds (6.17 mm/s vs 4.63 mm/s; *p*<0.001). This demonstrates that internal receptors (ACs) maintain static positional preferences while antennal inputs are essential for rapid boundary decisions, evidenced by an 82% reduction in U-turns after ablation (*p*<0.001).

These findings establish a new paradigm for understanding thermal navigation, where multiple sensory pathways operate in parallel with specialized functions. It can also assist when core locomotor rhythms (staccato pattern) remain invariant to thermal perturbations and hierarchical organization separates persistent positional maintenance (AC-dependent) from dynamic boundary responses (antennae-dependent). The work provides quantitative foundations for investigating how discrete sensory modalities integrate to produce adaptive navigation, with particular implications for understanding neural circuit organization in sensorimotor transformation. The precise behavioral measurements (avoidance indices, movement speeds, turn frequencies, centrophobism, pairwise encounters, and staccato manner) establish new standards for characterizing thermotactic behavior in genetic models.

## Methods

This study utilizes a two-choice behavioral assay to investigate temperature preference and locomotor activity in Drosophila melanogaster. The experimental setup consists of four Peltier modules arranged in a two-by-two configuration, integrated with thermal sensors and cooling mechanisms to create distinct thermal zones. To accurately model and control temperature dynamics, system identification experiments were conducted, applying various input currents and analyzing thermal responses under both active cooling and warming conditions. A proportional-integral-derivative (PID) control system was implemented to refine temperature regulation, ensuring stable and precise environmental conditions. Additionally, finite element method simulations were performed to characterize the thermal behavior of the system and assess the effects of cooling mechanisms. Locomotor activity was analyzed through a detection-based multi-object tracking framework, leveraging deep-learning-based object detection and trajectory reconstruction. To investigate the role of antennal thermal inputs in temperature-driven behavior, a subset of flies underwent surgical ablation of the final antennal segment, including the arista. Detailed descriptions of the experimental setup, control strategies, and data analysis methods are provided in the following sections:

### Components of the Behavioral Assay Device

The two-choice behavioral essay setup used in this study consists of four Peltier modules, each adhered to a fin using thermal adhesive. The assembly is mounted on a frame constructed from aluminum profiles and Plexiglass plates. Thermal sensors are adhered to the corners of each Peltier unit, which are organized in a 2×2 configuration (two rows and two columns). A fan is positioned beneath the fins and within the frame for cooling. More details about the structure are provided in the supplementary materials.

### System Identification and Control

Before conducting system identification, the setup operates as a black box, with electric current supplied to the Peltier module as the input and the sensed temperature as the output. System identification methods establish a relationship between these variables, enabling the development of a mathematical model for the system. System identification was performed under two conditions: cooling with the fan activated and warming without active cooling. Given the maximum allowable current of 2 amperes for the controller unit, input currents of 0.5, 1, and 1.5 amperes were applied, and the corresponding output temperatures were recorded. To prevent potential heat damage, the 1.5-ampere input was excluded from the warming experiments, as it led to rapid and excessive temperature increases. The system model was identified using a 1-ampere input and validated against the responses at other current levels. Additional details are provided in the supplementary materials.

#### Cooling and Heating Phases

For the cooling phase, the most accurate model capturing the undershoot in the system response consisted of two poles and one zero. As shown in *Figure 6 – a, b* the experimental and predicted outputs for a 1-ampere input exhibit strong agreement. However, model evaluation with 0.5 A and 1.5 A inputs yielded discrepancies. The predicted response for 0.5 A was positioned higher than the corresponding experimental result, whereas the predicted response for 1.5 A was lower. Following the same approach as in the cooling phase, system identification was performed using a 1-ampere input for heating. The two-pole, one-zero model showed promising results and fitted well to the experimental data, but did not perform well on the test current. These deviations arise from disturbances whose magnitudes do not scale linearly with the input current.

**Figure 6.**
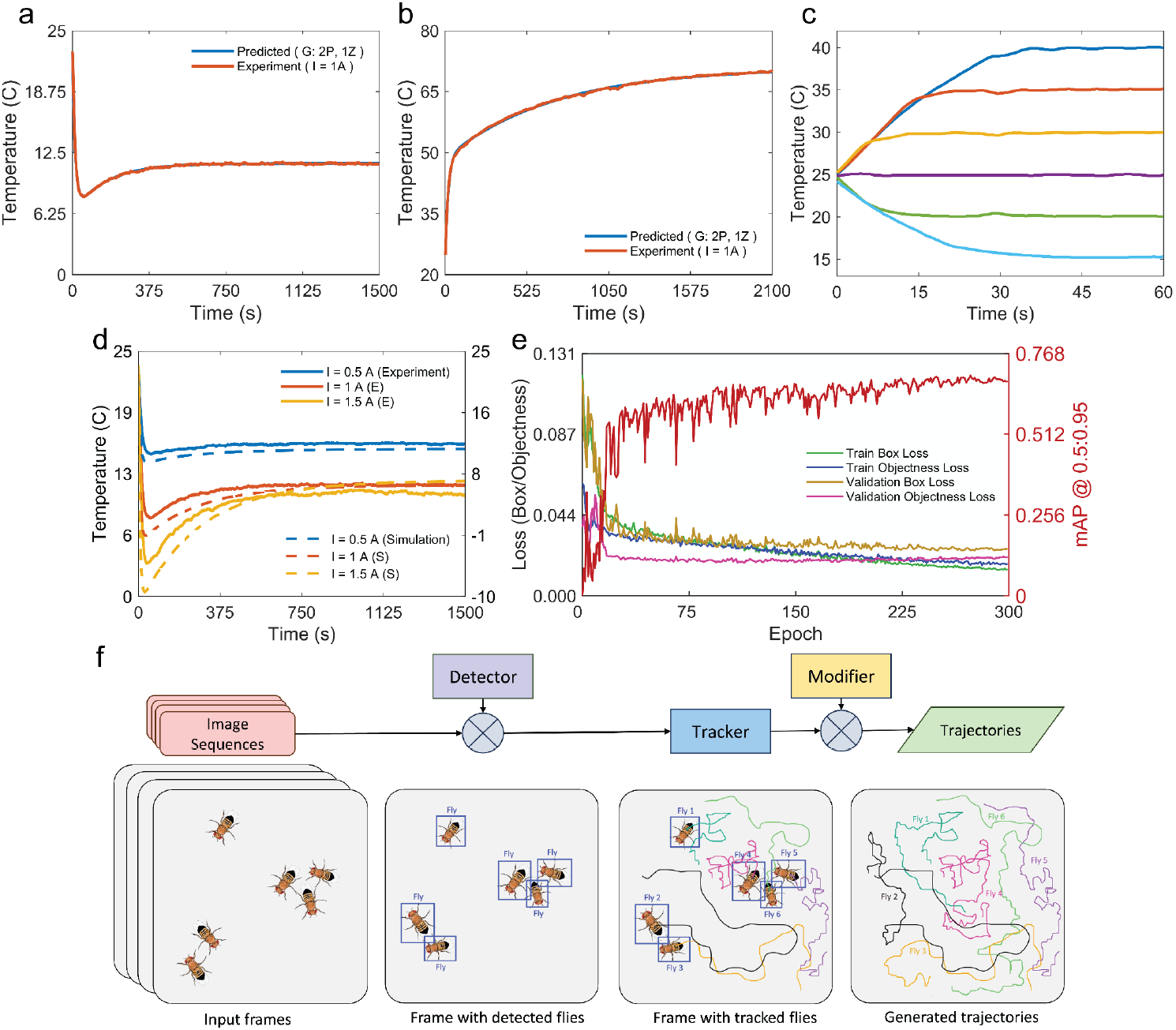
System Identification, Temperature Control, and Tracking Framework for Thermal Prefereance Assays. The initial temperature was approximately 23.9°C (mean) with a standard deviation of 1°C for the experiments. In panels (a-d), for the sake of consistancy in plots, some data were clipped to common threshholds. (a, b) System identification for heating and cooling was performed using temperature–time responses from a 1-A current applied to a Peltier module. A two-pole-one-zero model fit both processes. Model predictions were evaluated at 0.5 A and 1.5 A (not shown in the figures); for heating, 1.5 A was omitted to avoid sensor damage. Models failed to generalize beyond trained current levels, as discussed in the text. (c) Temperature control was achieved via PID algorithms with set points of 15–40°C. Rising time variations were due to current limits and PID parameters. A 0– 1.5°C offset was added to each set point during the control process when PID tuning alone could not ensure steady-state accuracy. (d) Comparison between experimental and simulated cooling results. Solid lines on the left indicate measured data; those on the right show simulations. While peak times matched well, steady-state temperatures diverged. (e) YOLOv5 model performance metrics—including bounding box regression and objectness loss—for both training and validation datasets are presented. (f) Overview of the detection-based multiple-fly tracking framework: (1) Image sequences are extracted from an offline video file; (2) The trained YOLOv5 fly detection model identifies individual flies; (3) The DeepSORT associates detections across frames. (4) Individual movement trajectories are reconstructed with correction applied to ensure accuracy.

#### PID Temperature Control

A microcontroller board and H-bridge modules were used to regulate the magnitude and direction of the current supplied to each Peltier module. The required current magnitude for temperature correction was determined using a PID controller, with its parameters adjusted through trial and error. Results demonstrated that while system identification provided valuable insights into the system dynamics, achieving precise temperature control required further refinement. To complement the identified model, a PID controller was implemented and tuned through an iterative process. Initially, all PID parameters were set to zero. The proportional gain was gradually increased to bring the system response closer to the desired set point. The integral gain was then adjusted to eliminate steady-state error while minimizing excessive oscillations. Finally, the derivative term was introduced to reduce the amplitude of oscillations and improve transient response. The control strategy aimed to achieve a peak time below 30 seconds and oscillation amplitudes within 0.5°C. Given the differing system dynamics in heating and cooling phases, distinct PID parameters were optimized for each case to enhance performance.

### FEM Simulation

Finite element simulations were performed to better understand the thermal dynamics of the system and assess the impact of cooling mechanisms, such as a fan, in preventing the system from reaching critical temperatures. These simulations serve as a complementary tool to experimental validation, offering insights into system behavior under varying conditions and helping to predict potential outcomes that may not be easily achievable through experiments alone. Given the significance of thermal regulation for Drosophila melanogaster experiments, the simulations were performed specifically for the cooling phase, where active cooling is critical. Current values of 0.5, 1, and 1.5 A were applied to the Peltier module-heatsink model, and the average temperature on the upper surface of the Peltier module was recorded. The spatial temperature distribution across each module exhibited a high degree of uniformity, with a maximum variation of only 0.5°C. This consistency confirms that the average surface temperature is a reliable parameter for assessing thermal behavior. The experimental setup consists of four Peltier modules, each mounted on a heatsink. For finite element simulations, a single Peltier module on a heatsink was modeled using COMSOL Multiphysics. An electric current was applied as the input, and the corresponding thermal response was analyzed. Simulations were conducted under two conditions: without active cooling and with a fan positioned at the bottom. The input current was varied to evaluate the system’s thermal behavior. Additional details of the simulation are provided in the supplementary materials. Simulated peak time results closely matched experimental observations. However, discrepancies between experimental and modeled temperatures were noted, which can be attributed to similar factors discussed in the system identification section. A more detailed exploration of these discrepancies is provided in the discussion section.

### Preparation of Flies

#### Conditions

For the experiments, well-fed male flies aged 2–7 days were tested during the daytime. To distinguish males from females, physical markers such as smaller body size, a distinct black stripe at the end of the abdomen, a more rounded abdomen, and the presence of sex combs were used. Flies were maintained on a diet consisting of flour, yeast, vinegar, sugar, agar, propionic acid, and banana. To sort and transfer flies to the arena, cooling anesthesia was used, a simple and cost-effective method that avoids long-term effects if excessive cooling is prevented. Flies were placed in a refrigerator for approximately 20 minutes, as prolonged exposure to extreme cold can be lethal. Carbon dioxide can also be used for anesthesia, but its effects tend to persist longer and are not recommended for these experiments.

#### Surgical Ablation of Antennal Thermal Inputs

This study also examines the locomotor behavior of antennae-ablated flies in response to noxious warming. Each antenna consists of three segments: the scape, the pedicel, and the funiculus, with the latter carrying the feather-like arista. Previous research on Drosophila temperature sensation has identified warm thermoreceptor cells within the arista, suggesting its role in thermal perception. To assess the influence of antennae on behavioral responses to heat, the final antennal segment, including the arista, was completely removed. Surgical ablation was performed on the first day after eclosion while anesthetized flies were kept in indirect contact with ice to minimize stress. The procedure was completed as quickly as possible to prevent adverse effects from prolonged cold exposure. During the ablation procedure, a surgical hook was used to secure the fly by its wings to prevent movement. Straight Jeweler’s forceps were used for ablation, and all procedures were conducted under an Olympus SZ3060 zoom stereo microscope. Flies were then transferred to food vials and allowed to recover for 1–2 days before testing.

### Data Acquisition from Two-choice Temperature Preference Assays

Experiments were conducted at room temperature. In each trial of the two-choice temperature preference assay, flies were presented with a choice between a base temperature (BT) of 25°C, an optimal temperature for Drosophila survival and reproduction, and a test temperature (TT) ranging from 25°C to 40°C in 5°C increments. Each trial lasted approximately two minutes, after which the spatial configuration of BT and TT quadrants was reversed for an additional two minutes. To ensure analysis focused on steady-state temperature conditions, the first 30 seconds of each trial were excluded from data processing. The interval between experiments was approximately 40 seconds, during which the device was turned off. A consistent sequence of test temperatures was used: (BT/TT) 25°/25°C, 25°/30°C, 30°/25°C, 25°/35°C, 35°/25°C, 25°/40°C, and 40°/25°C.

Locomotor behavior was recorded using a mobile phone camera at a frame rate of 60 FPS, later reduced to 30 FPS during editing with Adobe Premiere Pro to balance motion capture accuracy and file size efficiency (32 MB per video). A 1000 × 1000-pixel region of interest was extracted and saved for further analysis. The recorded data was then processed using a detection-based multiple-object tracking pipeline in Python to determine the position of each fly in each frame.

### Quantification of Locomotor Behavior via Video Tracking

A detection-based tracking (DBT) approach was implemented to develop a multi-fly tracking system, following a widely adopted strategy in modern multiple-object tracking (MOT) frameworks. In this method, the YOLOv5 network serves as an object detector, identifying and locating flies within each frame. The detected images are then processed by the DeepSORT algorithm, which assigns and maintains unique identities across successive frames, enabling the reconstruction of individual trajectories. Following the automated tracking process, potential errors were identified and corrected to improve accuracy. A detailed description of the complete tracking pipeline, including the training process for the YOLOv5 object detection model, DeepSORT implementation, and error correction procedures, is provided in supporting materials. The YOLOv5 loss function, optimized during the training process, consists of three distinct components: localization loss (also referred to as bounding box regression loss), which is responsible for object localization; confidence loss (objectness loss), which determines the presence of an object; and classification loss, which ensures category prediction accuracy. *Figure 6 – e* illustrates the localization and objectness losses on both the training and validation datasets, along with the mean Average Precision (mAP)@ [0.5, 0.95] for the top-trained model after 300 epochs. Given that only a single class is predicted, classification loss is zero and is therefore not shown. Evaluations indicate that, among the six trained models, the YOLOv5m model with a low-augmentation hyperparameter configuration exhibits the top performance. This model achieved the mAP@ [0.5, 0.95] of 69.8% on the training dataset. When applied to the validation and test datasets, with a confidence threshold of 0.7, the results were 72.0% and 69.7%, respectively. Complete results for the other trained models are available in the supplementary materials.

### Locomotor Feature Extraction

Multiple parameters are computed to construct the portrait of locomotor behavior: Avoidance Index (AI), the total traveled distance (D), the distance traveled in each test (D_test_), base (D_base_), and boundary (D_boundary_) regions, the number of active episodes (N_a_), the frequency ratio 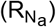 and length ratio 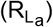 of active episodes, the average peak speed of the active episodes (V_peak_), the mean walking speeds in each test (V_test_), base (V_base_), and boundary regions (V_boundary_), the average relative area covered by the group (R_A_), the temporal ratio spent in each test (R_test_), base (R_base_), and boundary regions (R_boundary_), the temporal ratio spent in the peripheral area (R_p_), number index or frequency 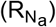 and length index 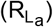 of pairwise encounters, the ratio of the U-turn (R_U-turn_). This description of locomotor behavior represents a tool which will facilitate the study of the role of the different stimulants in the organization of locomotor activity.

The positions were obtained by the explained tracking pipeline. In order to convert pixel-unit into real world mm-unit we applied conversion factor ≈ 0.053 (almost equal to 0.053), based on the radius of the chamber in the real world and its value in the frame. Along with position, “orientation angle” and “velocity” are also calculated as follows:

#### Orientation Angle

The general idea to determine the orientation angle of each fly in each frame of the video is to separate the fly from the background in each detected bounding box, using Otsu’s method for image thresholding accompanied by common morphological operations based on dilation and erosion in MATLAB, and then fit an ellipse to its body, using Principal Component Analysis (PCA) method. Thus, the angle that the major axis of the ellipse makes with the positive direction of the horizontal axis is considered as the angle of the fly. To get the angle of rotation of the fly between two consecutive frames, it is enough to subtract the orientation angle of the fly in these two specific frames.

#### Velocity

The velocity in both the horizontal and vertical direction can be calculated by taking the first order numerical derivative with respect to time, from the position in the same direction. But if this method is used, it should be ensured that the path is well smoothed in that specific direction. Fly’s zigzag movement and inaccurate trajectory tracking are among the causes of breaks in movement paths (sudden changes in direction). On the one hand, the presence of these breaks in the motion curve creates problems in its continuity and on the other hand, the undefinition of the noise derivative can lead to large errors and, as a result, meaningless velocity estimates. The solution is to use a curve fitting technique to smooth the time series. This methodology involves moving a window of a certain width (equal to 9 frames) across the data at a stride equal to the window’s width. The curve fitting process is performed inside each window. As the border point of each of the two successive windows is fitted with a different value, we take the average of those two values to be the output of that fitting. To ensure smoothness at the border points, we use the spline smoothing method (with its parameter equal to 0.995). For curve fitting inside each window, three smoothing options are available: spline, linear, and polynomials with the best fit according to the R-square criterion, which can be used depending on the volume of calculations. We found that the spline smoothing method with a parameter of 0.4 had the best performance (besides having a reasonable computational cost). This approach has been used throughout this research whenever curve fitting is mentioned. Finally, for calculating velocity, we used the central finite difference method with an 8th-order accuracy. It should be noted that for the first and last 4 points, we employed the forward and backward finite difference methods with a 6th-order accuracy, respectively. Moreover, the time interval between two successive frames is equal to the inverse of the camera frame rate, i.e., ∼0.033 sec.

Based on the positions of flies in each frame, multiple attributes computed. The Avoidance Index (AI) represents a simple measure of how much thermotaxis behavior is induced by the hot test temperatures. AI is calculated by differences in time spent in base and test regions and normalized by the entire experiment duration. In group assays, this property can be interpreted by the normalized difference of the number of flies in two regions of base and test temperature. Moreover, observations indicate that the test temperature influences the fly’s decision-making at the boundary. This decision is therefore categorized into two groups: cross and U-turn. In cross-motion, the fly traverses the base, boundary, and test regions sequentially. In contrast, the fly makes a U-turn decision when it leaves the base region, moves to the boundary region, and then returns to the original base region.

Having the speed in each frame, the staccato manner, a characteristic of fly’s locomotive behavior could be investigated. Previous studies ^43,45^ indicated that the locomotor behavior of flies, analogous to many other animals, is typically segmented into two (or more) motion regions. In other words, flies typically walk in a staccato manner, marked by bouts of fast walking episodes interspersed among stops or slowed speed. In the fly literature, these segments of differing speed are typically referred to as activity and inactivity and are defined by setting hard thresholds on the movement. Inactive episodes are defined as intervals where the speed falls below a threshold. This threshold was chosen 1 mm/sec by noting the maximum measured speed in segments of the video where the fly remained stationary to the eye of an observer. Thus, we can identify the sequence of active and inactive episodes within each video. Moreover, multiple related parameters, including the count (N_a_) and duration of both types of episodes, the ratio of active to inactive episode lengths 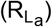, and the ratio of active to inactive episode counts 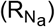, can be established. Furthermore, the maximum speed attained during each active episode can be determined, and the average of these maximum speeds can be calculated as the average peak speed of the active episodes (V_peak_).

In addition to the aforementioned factors, in each of the three areas—base region, boundary region, and test region—parameters such as average speed of activity (V_base_, V_boundary_, V_test_), temporal ratio of activity (R_base_, R_boundary_, R_test_), and traveled distance (D_base_, D_boundary_, D_test_) were also calculated. Furthermore, we took into account the average distance of flies from the chamber wall as a peripheral factor, so the temporal ratio spent in the peripheral area (from 0.65R to R) was calculated as a centrophobism behavior criterion (R_p_). Additionally, group parameters considered, including the average area covered by the group of six flies and pairwise encounters. The average area covered by the group (R_A_) indicates the dispersion of flies in the chamber. In each frame, the arrangement of 6 flies forms a convex polygon. The mean area of these polygons across successive frames is regarded as the average covered area in each experiment. Additionally, each pairwise encounter is considered in such a way that the distance between the centers of two flies is less than 95% of a fly’s body length, approximately 2.65 mm. We quantify this parameter for each video by calculating two ratios: 1) the ratio of the frequency of pairwise encounters to the total possible cases for a pairwise encounter (15 in total, 6 combinations of 2 without repetition), noted with 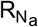, and 2) the ratio of the cumulative duration of encounters to the entire duration of the experiment 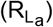.

### Statistical Analyses

Using statistical hypothesis tests, different groups were compared based on temperature and antennae status. For comparing multiple groups, there are different statistical methods depending on the number of groups, the type of data, and the research hypothesis, and parametric methods are preferred ^67-69^. Thus, according to the appropriate assumption of observations independency, Kolmogorov-Smirnov test was used to confirm normality of distribution and Levene’s test was used to confirm homogeneity of variance. One-Way ANOVA or Welch ANOVA tests have been used to compare wild-type flies at different temperatures, depending on whether these conditions are met. Also, at 35°C, wild and antennae-ablated fly groups were compared with independent sample t-tests with equal or unequal variances. For data that are not normally distributed, the non-parametric tests, Mann-Whitney U and Kruskal-Wallis, were employed as alternatives to the two-sample t-test and One-Way ANOVA, respectively. In all cases, the confidence interval for statistical tests is equal to 95% (5% significance level).

### Classification Methods

Classification is defined as a data mining (machine learning) technique to predict group membership for data instances ^70^. There are different types of classification algorithms, depending on the nature and complexity of the data and the problem. Some of the common classification algorithms are Decision Tree, Logistic Regression, Naïve Bayes (NB), Support Vector Machine (SVM), Artificial Neural Networks (ANN), K-Nearest Neighbor (KNN) and Ensemble Classifier ^71,72^. By utilizing the extracted features, this section investigates two problems: 1) Classifying four different temperature groups (25°C, 30°C, 35°C, 40°C) in the wild group, and 2) Classifying two groups of wild and antennae-ablated flies at 35C. The classification methods outlined above are applied to each of these two problems. The hyperparameter values were tuned by using Bayesian optimization for 100 iterations. As a criterion for choosing the best classifier, we looked at its classification accuracy on validation dataset. In order to validate, we used a 5-fold cross-validation method. Also, in assessing how the features contribute to determining the test temperature or antenna status, employing P values in one-way ANOVA provides a means to rank these features according to their impact.

### Mimicking Heat-Avoidance Behavior via a Braitenberg Vehicle Representation

A Braitenberg vehicle ^65^, an autonomous agent responding to sensor inputs, features primitive sensors measuring stimuli and wheels driven by motors serving as actuators. The interconnection of sensors and wheels leads to diverse behaviors, each with potential goal-oriented outcomes. The vehicle adapts its movement to attain certain situations and avoid others, showcasing a dynamic response to changing circumstances.

Critical to Braitenberg vehicles are sensorimotor couplings—direct connections between sensors and motors (*Figure 4 – a*). The nature and mechanism of these couplings determine the exhibited behavior. Ipsilateral couplings connect sensors and motors on the same side, while contralateral couplings link sensors on one side to motors on the opposite side ^66^. Couplings can be inhibitory or excitatory, with inhibitory relationships implying a weaker motor excitation with a stronger sensor stimulus and excitatory connections signaling a stronger motor excitation with a more intense sensor stimulus ^66^. This nuanced wiring system enables the vehicle to showcase a spectrum of behaviors, providing valuable insights into the complexity arising from simple designs in the realm of autonomous agents.

Based on the provided explanations, the heat avoidance behavior of fruit fly can be reproduced by a relatively straightforward “Braitenberg vehicle.” This model involves the fly, represented as the vehicle, actively avoiding a harmful heat source by utilizing the input from two symmetrical sensors and regulating the speed of their respective motors.

Towards this aim, the in-silico vehicle is scaled to match the fly’s physical dimensions, with motor distance equivalent to body width (1 mm, derived from the minor diameter of the ellipse in the segmentation method). The dynamic motion model aligns with the principles outlined in the influential paper ^39^:

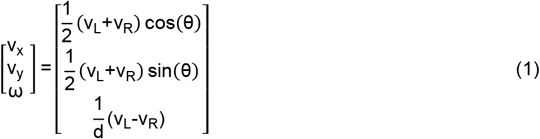

Where (v_x_, v_y_) is the velocity of the centroid, ω is the angular velocity, and θ is the orientation. Here, v_L_ and v_R_ are the velocities of the left and right motor, respectively, while d is the distance between the two wheels (set to 1 mm, consistent to a fly’s width). Sensory inputs are donated as s_L_ and s_R_ in which (s_L0_, s_R0_) is the perceived temperature by the left and right antennae that a normal random noise (ε_L_, ε_R_), a noise from the normal distribution with mean zero and standard deviation parameter 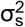 is also applied for each of them. We considered s_L0_=s_R0_=s_0_ that s_0_ equals to the temperature of the region (test region, base region, or boundary region) where the fly is standing in.

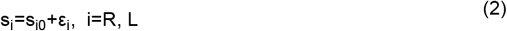

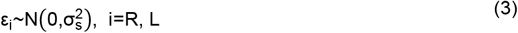

(ε_L_for the left sensor and ε_R_ for the right sensor).

The velocities of the wheels are expressed as linear combinations of sensory input (s_L_, s_R_), which undergo processing through a logistic nonlinearity (h) at the two symmetric sensors and are additively combined with the normal random noise γ and the initial velocity v_0_. w_I_ is ipsilateral weight and w_C_ is contralateral weight and the initial velocity v_0_ is set based on the experimental results:

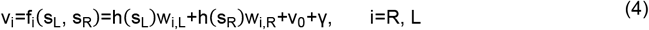

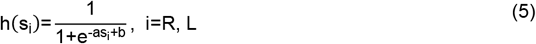

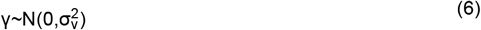

Note that the vehicle wiring is symmetric, so:

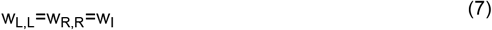

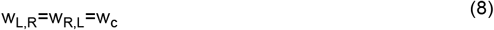

At time frame j, upon collision with the wall of the simulated chamber, the vehicle’s position was set to the rebound point and its velocity updated to 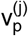. The position was then recalculated using the constant-velocity motion equation with 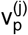.

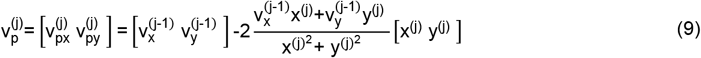

Additionally, based on the above description, parameters (a, b, w_I_, w_C_) must be fine-tuned to simulate the behavior of a fruit fly’s thermotaxis. Optimizing this vehicle could take into account objectives such as the avoidance index and the turn-cross ratio. Therefore, the general approach involves solving an optimization problem over a four-dimensional space and objective functions such as the ones mentioned earlier. This situation calls for multi-objective optimization methods such as the Non-dominated Sorting Genetic Algorithm II. However, performing an optimization process at this level is beyond the scope of this research. So, we will apply the trial-and-error method to determine values relatively close to the optimum value. It is important to emphasize that these values are not necessarily optimal, and a multi-objective optimization procedure is necessary to achieve a more accurate model.

Our strategy to specify values of ipsilateral (w_I_) and contralateral weights (w_C_) is region-based. It is also assumed that the sensor’s nonlinearity parameters (gain a and bias b) are constant (region-free). The general idea is that when a vehicle crosses from the base region into the boundary region, it is prevented from entering the test region through what we call an inhibitory action. In other words, with a suitable positive w_I_ (excitatory) and a suitable negative w_C_(inhibitory), we can guarantee the U-turn action at the boundary so that the vehicle returns to the same quarter of the base. However, if the current boundary position is close to the opposite base quadrant, its entry into that quadrant is strengthened. On the other hand, if the vehicle has crossed from the test region into the boundary region, it is encouraged to enter the base region as swiftly as possible using appropriate positive coefficients.

## Acknowledgements

We would like to express our gratitude to Amir Tarizadeh for his expert guidance on fly experimentation, to Mohammad Amin Mirzaee for his technical assistance with the temperature control apparatus, to Ramtin Tabatabaei for sharing his Peltier module model that was later used in this study and to Dr. Reza Naderloo for granting us access to the microscopes at the Laboratory of Biosystematics and Ecology at the University of Tehran, as well as his students for their assistance in capturing microscopic images.

## Data Availability

The data needed to reproduce the results is archived in Zenodo at https://doi.org/10.5281/zenodo.16739555. The behavioral analysis code can be found at https://github.com/hizeinab/multi-fly-thermotaxis-analysis.git.

## Additional Information

## Supporting Materials File

Details about the experimental setup, multiple-fly tracking pipeline and parameters of the Braitenberg vehicle are provided in the supporting materials PDF file.

## Author Contributions

- Conceptualization: ZM, SJ, EH
- System setup for the temperature control device: SJ
- Experimental design and performance: ZM, SJ
- Quantification of locomotor behavior: ZM
- Data analysis: ZM
- Visualization: ZM, SJ
- Manuscript preparation: ZM, SJ
- Supervision: EH, AS, TM

## References

1. Luo, L., Gershow, M., Rosenzweig, M., Kang, K., Fang-Yen, C., Garrity, P.A., and Samuel, A.D. (2010). Navigational decision making in Drosophila thermotaxis. Journal of Neuroscience 30, 4261–4272.

2. Tinette, S., Zhang, L., and Robichon, A. (2004). Cooperation between Drosophila flies in searching behavior. Genes, Brain and Behavior 3, 39–50.

3. Adams, M.D., Celniker, S.E., Holt, R.A., Evans, C.A., Gocayne, J.D., Amanatides, P.G., Scherer, S.E., Li, P.W., Hoskins, R.A., and Galle, R.F. (2000). The genome sequence of Drosophila melanogaster. Science 287, 2185–2195.

4. Iliadi, K.G. (2009). The genetic basis of emotional behavior: has the time come for a Drosophila model? Journal of neurogenetics 23, 136–146.

5. O’Kane, C.J. (2011). Drosophila as a model organism for the study of neuropsychiatric disorders. Molecular and functional models in neuropsychiatry, 37–60.

6. Pandey, U.B., and Nichols, C.D. (2011). Human disease models in Drosophila melanogaster and the role of the fly in therapeutic drug discovery. Pharmacological reviews 63, 411–436.

7. Kaun, K.R., Devineni, A.V., and Heberlein, U. (2012). Drosophila melanogaster as a model to study drug addiction. Human genetics 131, 959–975.

8. Kaufman, T.C. (2017). A short history and description of Drosophila melanogaster classical genetics: Chromosome aberrations, forward genetic screens, and the nature of mutations. Genetics 206, 665–689.

9. Bellen, H.J., Tong, C., and Tsuda, H. (2010). 100 years of Drosophila research and its impact on vertebrate neuroscience: a history lesson for the future. Nature Reviews Neuroscience 11, 514–522.

10. Dimitrijevic, N., Dzitoyeva, S., and Manev, H. (2004). An automated assay of the behavioral effects of cocaine injections in adult Drosophila. Journal of neuroscience methods 137, 181–184.

11. Chang, K.T., and Min, K.-T. (2005). Drosophila melanogaster homolog of Down syndrome critical region 1 is critical for mitochondrial function. Nature neuroscience 8, 1577–1585.

12. Bates, G.P., Dorsey, R., Gusella, J.F., Hayden, M.R., Kay, C., Leavitt, B.R., Nance, M., Ross, C.A., Scahill, R.I., and Wetzel, R. (2015). Huntington disease. Nature reviews Disease primers 1, 1–21.

13. McGurk, L., Berson, A., and Bonini, N.M. (2015). Drosophila as an in vivo model for human neurodegenerative disease. Genetics 201, 377–402.

14. Tian, Y., Zhang, Z.C., and Han, J. (2017). Drosophila studies on autism spectrum disorders. Neuroscience Bulletin 33, 737–746.

15. Harnish, J.M., Link, N., and Yamamoto, S. (2021). Drosophila as a model for infectious diseases. International journal of molecular sciences 22, 2724.

16. Layalle, S., They, L., Ourghani, S., Raoul, C., and Soustelle, L. (2021). Amyotrophic lateral sclerosis genes in Drosophila melanogaster. International Journal of Molecular Sciences 22, 904.

17. McDonald, J.M., Ghosh, S.M., Gascoigne, S.J., and Shingleton, A.W. (2018). Plasticity through canalization: the contrasting effect of temperature on trait size and growth in Drosophila. Frontiers in Cell and Developmental Biology 6, 156.

18. Soto-Padilla, A., Ruijsink, R., Span, M., van Rijn, H., and Billeter, J.-C. (2018). An automated method to determine the performance of Drosophila in response to temperature changes in space and time. JoVE (Journal of Visualized Experiments), e58350.

19. Truitt, A.M., Kapun, M., Kaur, R., and Miller, W.J. (2019). Wolbachia modifies thermal preference in Drosophila melanogaster. Environmental microbiology 21, 3259–3268.

20. Garrity, P.A., Goodman, M.B., Samuel, A.D., and Sengupta, P. (2010). Running hot and cold: behavioral strategies, neural circuits, and the molecular machinery for thermotaxis in C. elegans and Drosophila. Genes Dev 24, 2365–2382. 10.1101/gad.1953710.

21. Sayeed, O., and Benzer, S. (1996). Behavioral genetics of thermosensation and hygrosensation in Drosophila. Proceedings of the National Academy of Sciences 93, 6079–6084.

22. Zars, T. (2001). Two thermosensors in Drosophila have different behavioral functions. Journal of Comparative Physiology A 187, 235–242.

23. Barbagallo, B., and Garrity, P.A. (2015). Temperature sensation in Drosophila. Current opinion in neurobiology 34, 8–13.

24. Kafle, T., Grub, M., Sakagiannis, P., Nawrot, M.P., and Arguello, J.R. (2025). Evolution of temperature preference behaviour among Drosophila larvae. iScience 28.

25. Suito, T., and Tominaga, M. (2024). Functional relationship between peripheral thermosensation and behavioral thermoregulation. Frontiers in Neural Circuits 18, 1435757.

26. Capek, M., Arenas, O.M., Alpert, M.H., Zaharieva, E.E., Méndez-González, I.D., Simões, J.M., Gil, H., Acosta, A., Su, Y., and Para, A. (2025). Evolution of temperature preference in flies of the genus Drosophila. Nature, 1–9.

27. Gonzalez-Tokman, D., Cordoba-Aguilar, A., Dattilo, W., Lira-Noriega, A., Sanchez-Guillen, R.A., and Villalobos, F. (2020). Insect responses to heat: physiological mechanisms, evolution and ecological implications in a warming world. Biol Rev Camb Philos Soc 95, 802–821. 10.1111/brv.12588.

28. Hamada, F.N., Rosenzweig, M., Kang, K., Pulver, S.R., Ghezzi, A., Jegla, T.J., and Garrity, P.A. (2008). An internal thermal sensor controlling temperature preference in Drosophila. Nature 454, 217–220.

29. Gallio, M., Ofstad, T.A., Macpherson, L.J., Wang, J.W., and Zuker, C.S. (2011). The coding of temperature in the Drosophila brain. Cell 144, 614–624.

30. Frank, D.D., Jouandet, G.C., Kearney, P.J., Macpherson, L.J., and Gallio, M. (2015). Temperature representation in the Drosophila brain. Nature 519, 358–361.

31. Xiao, R., and Xu, X.Z.S. (2021). Temperature Sensation: From Molecular Thermosensors to Neural Circuits and Coding Principles. Annu Rev Physiol 83, 205–230. 10.1146/annurev-physiol-031220-095215.

32. Rooke, R., Rasool, A., Schneider, J., and Levine, J.D. (2020). Drosophila melanogaster behaviour changes in different social environments based on group size and density. Communications biology 3, 304.

33. Ramdya, P., Lichocki, P., Cruchet, S., Frisch, L., Tse, W., Floreano, D., and Benton, R. (2015). Mechanosensory interactions drive collective behaviour in Drosophila. Nature 519, 233–236.

34. Levine, J.D., Funes, P., Dowse, H.B., and Hall, J.C. (2002). Resetting the circadian clock by social experience in Drosophila melanogaster. Science 298, 2010–2012.

35. Li, K., and Gong, Z. (2017). Feeling Hot and Cold: Thermal Sensation in Drosophila. Neurosci Bull 33, 317–322. 10.1007/s12264-016-0087-9.

36. Dillon, M.E., Wang, G., Garrity, P.A., and Huey, R.B. (2009). Review: Thermal preference in Drosophila. J Therm Biol 34, 109–119. 10.1016/j.jtherbio.2008.11.007.

37. Yamada, Y., and Ohshima, Y. (2003). Distribution and movement of Caenorhabditis elegans on a thermal gradient. J Exp Biol 206, 2581–2593. 10.1242/jeb.00477.

38. Goodman, M.B., Klein, M., Lasse, S., Luo, L., Mori, I., Samuel, A., Sengupta, P., and Wang, D. (2014). Thermotaxis navigation behavior. WormBook, 1–10. 10.1895/wormbook.1.168.1.

39. Simões, J.M., Levy, J.I., Zaharieva, E.E., Vinson, L.T., Zhao, P., Alpert, M.H., Kath, W.L., Para, A., and Gallio, M. (2021). Robustness and plasticity in Drosophila heat avoidance. Nature communications 12, 2044.

40. Dell, A.I., Bender, J.A., Branson, K., Couzin, I.D., de Polavieja, G.G., Noldus, L.P., Pérez-Escudero, A., Perona, P., Straw, A.D., and Wikelski, M. (2014). Automated image-based tracking and its application in ecology. Trends in ecology & evolution 29, 417–428.

41. Egnor, S.R., and Branson, K. (2016). Computational analysis of behavior. Annual review of neuroscience 39, 217–236.

42. Risse, B., Mangan, M., Del Pero, L., and Webb, B. (2017). Visual tracking of small animals in cluttered natural environments using a freely moving camera. pp. 2840–2849.

43. Martin, J.-R. (2004). A portrait of locomotor behaviour in Drosophila determined by a video-tracking paradigm. Behavioural processes 67, 207–219.

44. Besson, M., and Martin, J.R. (2005). Centrophobism/thigmotaxis, a new role for the mushroom bodies in Drosophila. Journal of neurobiology 62, 386–396.

45. Valente, D., Golani, I., and Mitra, P.P. (2007). Analysis of the trajectory of Drosophila melanogaster in a circular open field arena. PloS one 2, e1083.

46. Branson, K., Robie, A.A., Bender, J., Perona, P., and Dickinson, M.H. (2009). High-throughput ethomics in large groups of Drosophila. Nature methods 6, 451–457.

47. Simon, J.C., and Dickinson, M.H. (2010). A new chamber for studying the behavior of Drosophila. Plos one 5, e8793.

48. Ofstad, T.A., Zuker, C.S., and Reiser, M.B. (2011). Visual place learning in Drosophila melanogaster. Nature 474, 204–207.

49. Giraldo, D., Adden, A., Kuhlemann, I., Gras, H., and Geurten, B.R. (2019). Correcting locomotion dependent observation biases in thermal preference of Drosophila. Scientific reports 9, 3974.

50. Khaira, H., Trinidad, T., Luu, J., Khan, S., and Crockenberg, N. (2019). Measuring the effects of larval life stress on adult anxiety behavior in Drosophila melanogaster. J. Biol. Sci 5, 6.

51. Sridhar, V.H., Roche, D.G., and Gingins, S. (2019). Tracktor: image-based automated tracking of animal movement and behaviour. Methods in Ecology and Evolution 10, 815–820.

52. Panadeiro, V., Rodriguez, A., Henry, J., Wlodkowic, D., and Andersson, M. (2021). A review of 28 free animal-tracking software applications: current features and limitations. Lab animal 50, 246–254.

53. Dowse, H.B., Ringo, J.M., and Barton, K.M. (1986). A model describing the kinetics of mating in Drosophila. Journal of theoretical biology 121, 173–183.

54. Dankert, H., Wang, L., Hoopfer, E.D., Anderson, D.J., and Perona, P. (2009). Automated monitoring and analysis of social behavior in Drosophila. Nat Methods 6, 297–303. 10.1038/nmeth.1310.

55. Van Breugel, F., and Dickinson, M.H. (2012). The visual control of landing and obstacle avoidance in the fruit fly Drosophila melanogaster. Journal of Experimental Biology 215, 1783–1798.

56. Hoyer, S.C., Eckart, A., Herrel, A., Zars, T., Fischer, S.A., Hardie, S.L., and Heisenberg, M. (2008). Octopamine in male aggression of Drosophila. Current Biology 18, 159–167.

57. Kabra, M., Robie, A.A., Rivera-Alba, M., Branson, S., and Branson, K. (2013). JAABA: interactive machine learning for automatic annotation of animal behavior. Nature methods 10, 64–67.

58. Eyjolfsdottir, E., Branson, S., Burgos-Artizzu, X.P., Hoopfer, E.D., Schor, J., Anderson, D.J., and Perona, P. (2014). Detecting social actions of fruit flies. (Springer), pp. 772–787.

59. Pavlou, H.J., and Goodwin, S.F. (2013). Courtship behavior in Drosophila melanogaster: towards a ‘courtship connectome’. Current opinion in neurobiology 23, 76–83.

60. Bellemer, A. (2015). Thermotaxis, circadian rhythms, and TRP channels in Drosophila. Temperature 2, 227–243.

61. Mohammad, F., Aryal, S., Ho, J., Stewart, J.C., Norman, N.A., Tan, T.L., Eisaka, A., and Claridge-Chang, A. (2016). Ancient anxiety pathways influence Drosophila defense behaviors. Current Biology 26, 981–986.

62. Zanini, D., Giraldo, D., Warren, B., Katana, R., Andrés, M., Reddy, S., Pauls, S., Schwedhelm-Domeyer, N., Geurten, B.R., and Göpfert, M.C. (2018). Proprioceptive opsin functions in Drosophila larval locomotion. Neuron 98, 67-74. e64.

63. Wu, S., Tan, K.J., Govindarajan, L.N., Stewart, J.C., Gu, L., Ho, J.W.H., Katarya, M., Wong, B.H., Tan, E.-K., and Li, D. (2019). Fully automated leg tracking of Drosophila neurodegeneration models reveals distinct conserved movement signatures. PLoS biology 17, e3000346.

64. Soibam, B., Mann, M., Liu, L., Tran, J., Lobaina, M., Kang, Y.Y., Gunaratne, G.H., Pletcher, S., and Roman, G. (2012). Open-field arena boundary is a primary object of exploration for Drosophila. Brain and behavior 2, 97–108.

65. Braitenberg, V. (1986). Vehicles: Experiments in synthetic psychology (MIT press).

66. Shaikh, D., and Rañó, I. (2020). Braitenberg vehicles as computational tools for research in neuroscience. Frontiers in bioengineering and biotechnology 8, 565963.

67. Whitley, E., and Ball, J. (2002). Statistics review 5: Comparison of means. Critical Care 6, 1–5.

68. Bewick, V., Cheek, L., and Ball, J. (2004). Statistics review 9: one-way analysis of variance. Critical care 8, 1–7.

69. Anderson, D.R., Sweeney, D.J., Williams, T.A., Camm, J.D., and Cochran, J.J. (2016). Statistics for business & economics (Cengage Learning).

70. Kesavaraj, G., and Sukumaran, S. (2013). A study on classification techniques in data mining. (IEEE), pp. 1–7.

71. Neelamegam, S., and Ramaraj, E. (2013). Classification algorithm in data mining: An overview. International Journal of P2P Network Trends and Technology (IJPTT) 4, 369–374.

72. Osisanwo, F., Akinsola, J., Awodele, O., Hinmikaiye, J., Olakanmi, O., and Akinjobi, J. (2017). Supervised machine learning algorithms: classification and comparison. International Journal of Computer Trends and Technology (IJCTT) 48, 128–138.

73. Redmon, J., Divvala, S., Girshick, R., and Farhadi, A. (2016). You only look once: Unified, real-time object detection. pp. 779–788.

74. Razzok, M., Badri, A., El Mourabit, I., Ruichek, Y., and Sahel, A. (2023). Pedestrian detection and tracking system based on Deep-SORT, YOLOv5, and new data association metrics. Information 14, 218.

75. Zoph, B., Cubuk, E.D., Ghiasi, G., Lin, T.-Y., Shlens, J., and Le, Q.V. (2020). Learning data augmentation strategies for object detection. (Springer), pp. 566–583.

76. Bell, S., Zitnick, C.L., Bala, K., and Girshick, R. (2016). Inside-outside net: Detecting objects in context with skip pooling and recurrent neural networks. pp. 2874–2883.

77. Wojke, N., Bewley, A., and Paulus, D. (2017). Simple online and realtime tracking with a deep association metric. (IEEE), pp. 3645–3649.

78. Bewley, A., Ge, Z., Ott, L., Ramos, F., and Upcroft, B. (2016). Simple online and realtime tracking. (Ieee), pp. 3464–3468.

79. Pereira, R., Carvalho, G., Garrote, L., and Nunes, U.J. (2022). Sort and deep-SORT based multi-object tracking for mobile robotics: Evaluation with new data association metrics. Applied Sciences 12, 1319.

80. Dosovitskiy, A., Beyer, L., Kolesnikov, A., Weissenborn, D., Zhai, X., Unterthiner, T., Dehghani, M., Minderer, M., Heigold, G., and Gelly, S. (2020). An image is worth 16×16 words: Transformers for image recognition at scale. arXiv preprint arXiv:2010.11929.

81. Radford, A., Kim, J.W., Hallacy, C., Ramesh, A., Goh, G., Agarwal, S., Sastry, G., Askell, A., Mishkin, P., and Clark, J. (2021). Learning transferable visual models from natural language supervision. (PMLR), pp. 8748–8763.

