## Supporting Materials File for "Thermotaxis behavior of *Drosophila melanogaster*: A quantitative analysis of sensory-motor integration and heat avoidance"

Zeinab Moradi *et al.*

The primary components of this research aimed at analyzing thermal preferences involve two key steps: (1) data acquisition through two-choice temperature preference assays with flies, and (2) quantification of their locomotor behavior. The experimental setup required selecting appropriate hardware for the physical arrangement, conducting system identification, programming the temperature control logic, and performing finite element analysis (FEA) simulations. To reconstruct individual fly trajectories, we implemented a detection-based multi-fly tracking pipeline. Specifically, the YOLOv5 model was trained to detect the flies, while the DeepSORT algorithm was employed to assign and maintain consistent identities across successive frames. Subsequently, potential sources of error were identified and corrected to enhance the accuracy of the tracking system. As the detailed procedures exceed the scope of the main text, they are provided in the supplementary information.

### Details of the Experimental Setup

**Temperature-Controlled Device for Two-choice Behavioral Assay**

The Arduino Uno is an open-source microcontroller board with 14 digital I/O pins, 6 PWM pins, and 6 analog input pins, making it a suitable choice for controlling four Peltier modules. Peltier modules act as heaters or coolers depending on the direction of the current flowing through them. Owing to the thermoelectric (Peltier) effect, when a voltage and current are applied, one side of a Peltier module absorbs heat (producing a cooling effect) while the opposite side dissipates heat. Model TEC1-12710 Peltier elements were selected for temperature regulation. To reverse the current direction, two L298 H-bridge driver modules were employed. Each L298, conventionally used for controlling the speed and direction of servomotors, can independently drive two modules. In the present setup, each Peltier element is addressed through three microcontroller pins: two digital pins to set the current direction and one PWM line to modulate the drive amplitude. Temperature feedback was provided by DS18B20 digital thermal sensors. The manufacturer specifies that the DS18B20 sensors provide an accuracy of ±0.5 °C within the range −10 °C to 85 °C. To verify this claim, each sensor was wrapped in aluminum foil and immersed in a melting ice–water bath; the bath was continuously stirred to maintain a uniform temperature. The sensor readings were compared with those of a mercury thermometer. The procedure was repeated several times, and on every trial the difference between the thermometer and sensor readings remained below 0.5 °C.

During cooling operation, heat conducted from the hot face of a Peltier element can elevate the temperature of the cold face. To dissipate this heat, the original aluminum heat sink was cut into four independent sections, each ≈ 40 mm (h) × 45 mm (w) × 45 mm (l). Mounting all four modules on a single sink would have facilitated undesirable heat transfer between devices and, because the modules must be bonded to the sink, the failure of one unit would have rendered the entire assembly unusable. Bonding each module to its own heat sink therefore proved advantageous.

Because passive heat sinking alone was insufficient, a 90 mm × 90 mm × 25 mm axial fan (4.56 W) was installed beneath the sinks to enhance convective cooling. The Peltier elements were attached to their respective sinks—and the DS18B20 sensors to the module surfaces—using a thermally conductive adhesive. This adhesive minimizes interfacial thermal resistance; without it, the modules cool more slowly and the sensors report inaccurate temperatures.

The chamber walls were fabricated from white acrylic (plexiglass) and the roof from glass. The enclosure was modelled in SolidWorks and laser-cut from the acrylic sheet. In unmodified arenas, Drosophila readily climb the side walls and traverse the ceiling, thereby confounding vision-based tracking algorithms and exposing the insects to temperatures that differ from those at the floor. To mitigate these issues, we implemented three measures. First, we incorporated a zig-zag pattern into the interior walls, following the approach outlined in [39], to deter climbing behavior. Second, the chamber height was reduced to 1.5 mm, effectively restricting the flies' ability to transition freely between the floor and ceiling. It is important to note that the optimal chamber height may vary depending on the specific behavior being investigated [47]. Third, a transparent hydrophobic coating was applied to the glass ceiling, creating a sufficiently slippery surface to prevent the flies from walking on it. Alternatively, coatings such as Sigmacote® [47] or Fluon [54] may be used to achieve a similar effect. The final chamber dimensions were 55 mm × 55 mm, featuring a zig-zag circular opening with an inner diameter of 46 mm and an outer diameter of 50.42 mm.

A MEGATEK MP-3005D DC bench supply provided power to the thermoelectric circuit. The unit offers two independent input–output channels (each input paired with one output) with adjustable voltage and current; each channel was assigned to two Peltier modules. Maximum voltage and current settings were limited to values safely within the ratings of all circuit components. During cooling operation, a second power supply was added to drive the fan.

The instrument frame was assembled from aluminum profiles. Ventilation patterns were designed in CAD and laser-cut into a plexiglass (acrylic) plate, which was mounted inside the frame. Four heat-sink/Peltier/sensor assemblies were arranged on this plate in a 2 × 2 configuration, while a fan was placed beneath the plate so that airflow could pass through the holes, over the fins, and dissipate waste heat. A laser-cut, cross-shaped acrylic barrier (1 mm thick) was inserted between the assemblies to minimize lateral heat conduction and to maintain fixed spacing; its height matched that of the assemblies. The top surfaces of the Peltier modules were covered with thin white paper tape to provide a flat, replaceable interface. When the tape was renewed, alignment marks were redrawn to ensure that the behavioral chamber remained centred. Illumination was supplied from above by two 9 W LED lamps, adjusted to produce a uniform light field across the arena.

**System Identification**

The MATLAB System Identification Toolbox was employed to characterize the system's thermal response. A step function served as the time-domain input signal; for cooling a negative sign was added behind the step function. The output was the temperature deviation from its initial value, sampled at 0.7260 s intervals. Transfer-function estimation was performed with the toolbox function tfest, which by default initializes parameters using the instrumental-variables (IV) method and then refines them via nonlinear least-squares optimization.

**Control**

Regulating a Peltier element requires control of both current direction and magnitude. Direction is set by the two logic inputs on the L298 H-bridge. When the inputs are identical (00 or 11), the bridge is disabled and no current flows; when they are complementary (01 or 10), current flows through the module in the corresponding direction. A negative temperature error (measured temperature < set-point) calls for a direction that heats the sensor region, whereas a positive error requires the opposite direction. Current amplitude is adjusted via the H-bridge's PWM control pin. The PWM duty cycle is calculated with a proportional–integral–derivative (PID) algorithm, and the PID gains were tuned empirically. If only a few discrete set-point temperatures are used, we recommend optimizing the PID parameters separately for each set-point to achieve tighter temperature regulation.

**FEM Simulation**

A single Peltier module mounted on a heat sink was simulated in COMSOL Multiphysics. The module comprises alternating n- and p-type semiconductor legs sandwiched between thin copper conductors. The CAD geometry, created in SolidWorks, was imported and assembled in COMSOL. Aluminum was assigned to the heat sink and alumina to the outer ceramic plates of the module. The semiconductor legs were modelled as bismuth telluride. To differentiate n-type from p-type elements, the Seebeck coefficient of half of the legs was given a negative sign.

The Thermoelectric Effect multiphysics interface was enabled in COMSOL, automatically coupling the Heat Transfer in Solids and Electric Currents physics. Domain assignments were then refined: every domain was included in Heat Transfer in Solids, while only the semiconductor legs and copper conductors were assigned to Electric Currents. The Thermoelectric Effect formulation itself was applied exclusively to the semiconductor domains.

In a typical Peltier module, most copper traces link an n-type leg to an adjacent p-type leg; however, two copper pads contact only a single leg—one n-type, the other p-type. One of these pads functions as the current input and the other as the terminal. The choice of which pad is driven positive and which is grounded sets the current direction through the couples, thereby determining which ceramic face becomes the hot side and which becomes the cold side.

To represent the forced-convection effect of the cooling fan, a Heat Flux boundary condition was imposed on the underside of the heat sink and on the internal faces between the fins. We assigned heat transfer coefficient of $25 W/m^2$ to this boundary condition, as it is the heat transfer coefficient for forced heat convection of the air. The rest of the faces were assumed to be exposed to air with the heat transfer of $1 W/m^2$.

The equations solved for the heat transfer physics are the energy conservation equation and Fourier's law of heat conduction, and are as the following:

$$\rho C_p \left( \frac{\partial T}{\partial t} \right) + \rho C_p u.\nabla T + \nabla.q = Q + Q_{ted} \tag{S 1}$$

$$q = -k\nabla T \tag{S 2}$$

Where ρ is density, $C_p$ is specific heat capacity at constant pressure, T is temperature, t is time, u is the velocity vector field, q is Heat flux vector, Q is external heat source per unit volume, $Q_{ted}$ is the rate of heat generation per volume, and k is thermal conductivity of the material. The dependent variable is Temperature. Quadratic Lagrange basis function is used for discretization. For the electric current physics, the following equations are solved:

$$\nabla.J = Q_{j,v} \tag{S 3}$$

$$J = \sigma E + \frac{\partial D}{\partial t} + J_e \tag{S 4}$$

$$E = -\nabla V \tag{S 5}$$

In which J is the current density vector, $Q_{j,v}$ is volume charge density, σ is electrical conductivity, E is electric field vector, D is electrical displacement vector, $J_e$ is external current density and V is electric potential. The basis function and dependent variable in this Physics are Quadratic and electric potential respectively.

For the Multiphysics part, two physics of "heat transfer in solids" and "electric current" are coupled together, and that includes two effects; thermoelectric effect and electromagnetic heating. For the thermoelectric effect, the following holds:

$$P=ST \tag{S 6}$$

$$q=PJ \tag{S 7}$$

$$J_e = -\sigma S \nabla T \tag{S 8}$$

Where P is power and S is Seebeck coefficient. And for the electromagnetic heating we will have the below equations:

$$\rho C_p \left( \frac{\partial T}{\partial t} \right) + \rho C_p u.\nabla T = \nabla.(k \nabla T) + Q_e \tag{S 9}$$

In which $Q_e$ is the heat source or sink term (external heat generation or absorption). The terms which are functions of time are omitted from the equations in the steady state studies.

### Multiple-fly Tracking Pipeline

The detection-based multi-fly tracking framework is structured in sequential steps. Initially, image sequences are extracted from an offline video file to provide the input for subsequent analyses. These sequences are then processed by a trained YOLOv5 object detection model, which identifies individual flies in each frame. Detected flies are subsequently tracked across consecutive frames using the DeepSORT algorithm, which associates and maintains individual identities throughout the sequence. This process enables the reconstruction of individual movement trajectories over time. In later sections, the procedures for detection, tracking, and error correction will be described in detail.

#### YOLOv5 Detection Model

The “You Only Look Once” (YOLO) algorithm is a renowned object detection method that partitions images into grids, with each grid cell tasked with object detection within its boundaries [73].The architecture of the YOLO family is simple, and its neural network can directly yield the location and class of the bounding box, facilitating real-time detection in videos. Owing to its speed and accuracy, YOLO has gained significant recognition in the field of object detection. By detecting objects using the entire image, YOLO can encode global information, thereby reducing the likelihood of erroneously detecting the background as an object. The fifth version of the YOLO algorithm called YOLOv5 was introduced in 2020 by Glenn Jocher, founder of Ultralytics, based on the PyTorch framework (For more details refer to the Ultralytics document at https://docs.ultralytics.com/yolov5/). YOLOv5 is categorized into five model versions based on network depth and width: YOLOv5n (nano), YOLOv5s (small), YOLOv5m (medium), YOLOv5l (large), and YOLOv5x (extra-large). Figure S 1 depicts the structure of YOLOv5. Here we investigated the performance of nano, small, and medium versions to detect flies in each frame of the video.

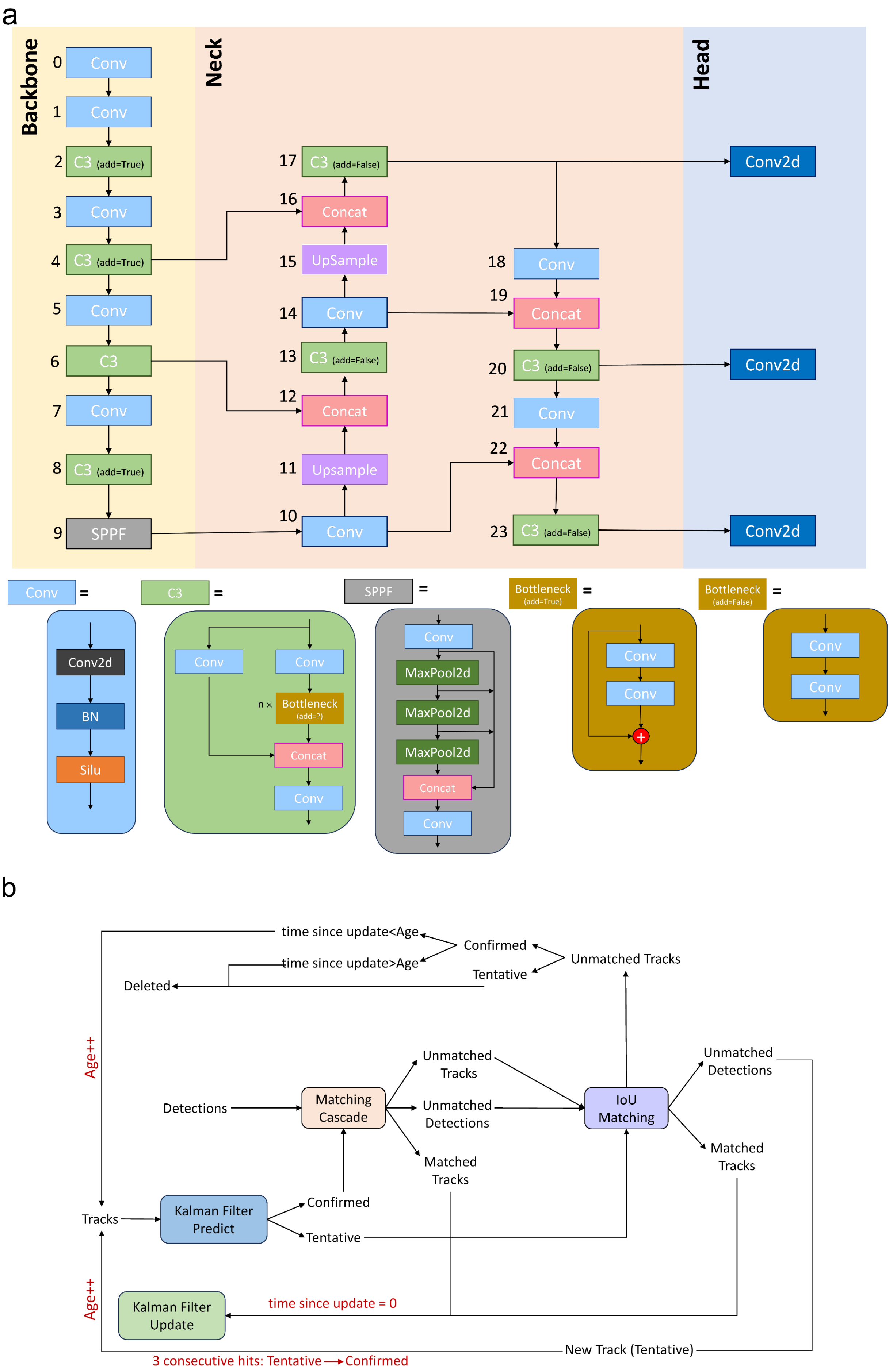


**Figure S 1. Detector and Tracker Architecture in the Fly Tracking Pipeline.** (a) YOLOv5 detection network is composed of three main components: Backbone, Neck, and Head. The Backbone extracts essential visual features using convolutional (Conv) layers, C3 modules, Bottleneck modules, and the Spatial Pyramid Pooling–Fast (SPPF) block. The Neck aggregates multi-scale feature maps to enhance object localization, while the Head generates final predictions, including bounding boxes, class probabilities, and objectness scores. For more information, please visit the Ultralytics YOLO documentation at https://docs.ultralytics.com/yolov5/. (b) The DeepSORT tracking algorithm integrates motion and appearance information to maintain object identity across frames. It uses Kalman filtering for motion prediction and a deep appearance descriptor for re-identification, enabling robust association of fly detections over time. The DeepSORT algorithm is overviewed in references [74].

***Training YOLOv5 Object Detection Model***: Multiple videos were taken of flies in the chamber and 160 photos were extracted from them. Bounding box labeling was done using LabelImg, a graphical python-based image annotation tool that enables exporting annotations directly in YOLO format. Afterwards, we divided the dataset into three training, validation, and test categories using a 70/20/10 split as a rule-of-thumb in splitting a dataset this size.

Data augmentation is a critical component of training deep learning models, which enables the models to generalize and perform better on unseen images. Considering the cost of annotating images for object detection, data augmentation may be even more crucial for this computer vision task [75]. So, data augmentation methods were applied to the training dataset in order to create 3 versions of each source image. Random rotation of between -15 and +15 degrees, Random Gaussian blur of between 0 and 1.5 pixels, Salt and pepper noise to 5 percent of pixels, and Mosaic are the methods used. Through augmentation, the dataset was expanded from 160 images to 384 images.

Nano, small, and medium versions of the YOLOv5 network were trained during 300 epochs, using an image input size of 640 × 640, a batch size of 32, and the default SGD optimizer. In addition, the weights were initialized using the COCO pre-trained model, and the hyperparameters were set using YAML-format configuration files (for low and medium augmentation) from the official repository. The performance of the models were evaluated based on mean Average Precision (mAP) averaged over IoU thresholds in [0.5:0.05:0.95] (simply denoted as mAP@[0.5:0.95] or mAP@[0.5, 0.95]). It is a common object detection metric, also used as standard metric of MS COCO dataset, places a significantly larger emphasis on localization accuracy [76].

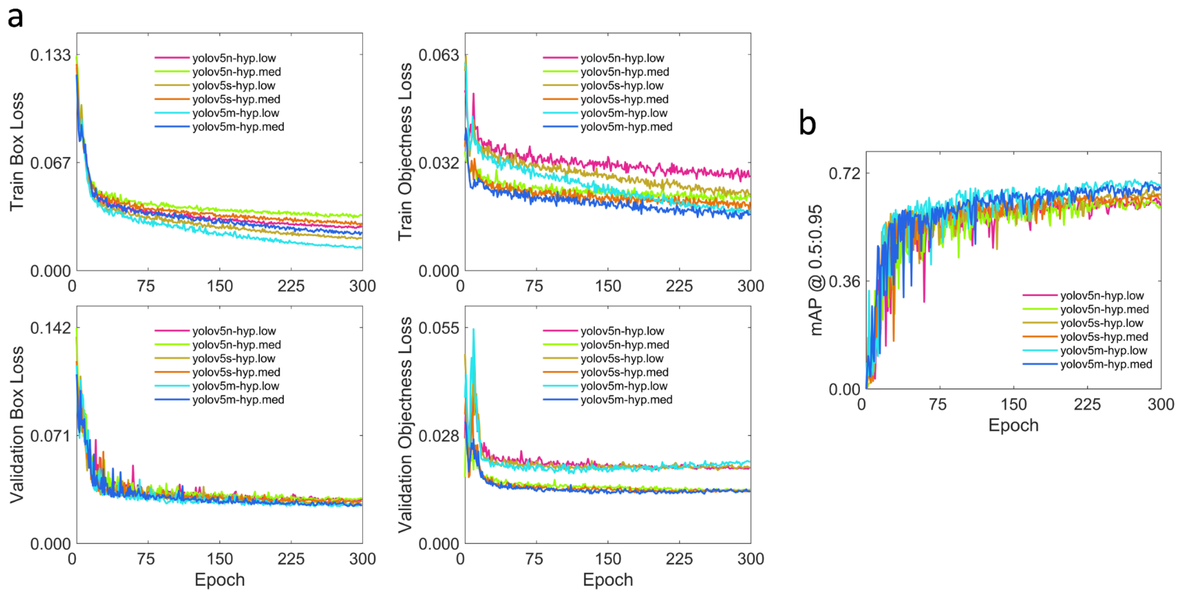


**Figure S 2. YOLOv5 Model Performance in Fly Detection Task.** YOLOv5m trained with a low-augmentation hyperparameter configuration achieved the best performance. (a) Localization loss and objectness loss curves on both training and validation datasets demonstrate stable convergence. (b) Mean Average Precision (mAP@[0.5, 0.95]) over training epochs indicates high detection accuracy. In real-world test samples, the trained model consistently identified flies with over 90% confidence and accurate localization.

### Deep-SORT Tracking Algorithm

The Deep-SORT algorithm [77], which leverages deep learning to improve object tracking accuracy and robustness in real-time video streams, is an advanced version of a simple and efficient online tracking algorithm called SORT [78]. While SORT struggles with tracking multiple objects that are closely located or occluded, Deep-SORT overcomes these challenges by using deep learning to associate detections of the same object across frames. The algorithm extracts the features from the object detection output and computes the similarity between detections. This allows Deep-SORT to accurately track multiple objects that are closely located or occluded, enabling the reidentification of tracks after a long period of occlusion. Deep learning also makes Deep-SORT more resilient to changes in appearance and lighting conditions, leading to more robust object tracking. The corresponding Deep-SORT modules resemble the Kalman Filter estimation and track management modules. An overview of the DeepSORT algorithm is provided in the works of [74,79]. Like SORT, the Hungarian algorithm uses a two-stage matching cascade to assign detected bounding boxes to tracks. In the first stage, the Deep-SORT method is used to match valid tracks based on motion and appearance metrics. The second stage, which uses the same data association strategy as SORT, links unpaired and tentative tracks with unpaired detections.

The integration of motion information involves computing the (squared) Mahalanobis distance between predicted states and detections. Along with this distance metric, a second metric that uses the smallest cosine distance measures the distance between each track and the appearance features of each measurement. Appearance features are typically computed using a pre-trained appearance descriptor network. In this case, we will need to re-label the collected dataset and train a second model in addition to the object detection model. Taking advantage of the idea described in Roboflow (For more details refer to Roboflow blog at https://blog.roboflow.com/zero-shot-object-tracking/, we used the efficient vision transformer model ViT-B/16 [80], trained as an image encoder in the robust, flexible, and highly generalizable CLIP model [81]. The pre-trained model is provided by the authors in their repository (https://github.com/OpenAI/CLIP). For the Deep-SORT, the following constant values were used: the hyperparameter $\lambda=0$ to control the influence of each metric on the combined association cost, the maximum age of 60, and a gating threshold of 0.4 for cosine distance metric. Moreover, all experiments were performed using the Tesla T4 GPU on Google Colaboratory.

### Modifying Possible Errors

The object tracking algorithm may exhibit suboptimal performance in certain frames, particularly in the assignment of unique identifiers. This can occur when the algorithm incorrectly assumes that a new fly has entered the scene, leading to the assignment of a new ID, or when an identity switch takes place. Another source of error stems from the failure of the object detector. This can manifest as a temporary loss of the fly's position in certain frames or as incorrect detections. Additional scenarios include instances when a fly exhibits an abrupt change in speed. In such cases, the constant-velocity motion model is inadequate for predicting the trajectories (e.g., when a fly jumps), or when, due to chasing behavior, a fly comes so close to another that they partially overlap. These inaccuracies can significantly impact the overall performance of the tracking system. So, it's crucial to address these issues to ensure the reliability and accuracy of the obtained trajectories.

We found that an unsuitable threshold value for the Non-Maximum Suppression (NMS) method in an object detection network can result in additional incorrect detections.

These incorrect detections are initially removed. Once all corrections related to detection errors have been applied and we have ensured that there are at most six detections (corresponding to the size of the fly group) in each frame, we proceed to corrections related to ID assignment and prediction of lost frames. In the following, the issue of erroneously assigning a new ID due to the incorrect assumption of a new object entering the scene is addressed. This is achieved by identifying the previous detection that has the smallest Euclidean distance to the current measurement. Moreover, ID switch errors are manually corrected by visually monitoring the trajectories. For lost data, the speed in the existing frame is estimated using a 4th-order finite difference approximation of the first derivative, and the prediction is made using the constant speed model. The predicted value is then used to replace the lost position.

### Spatial Maps of Temperature Preference for Test Temperatures on Quadrants I and III

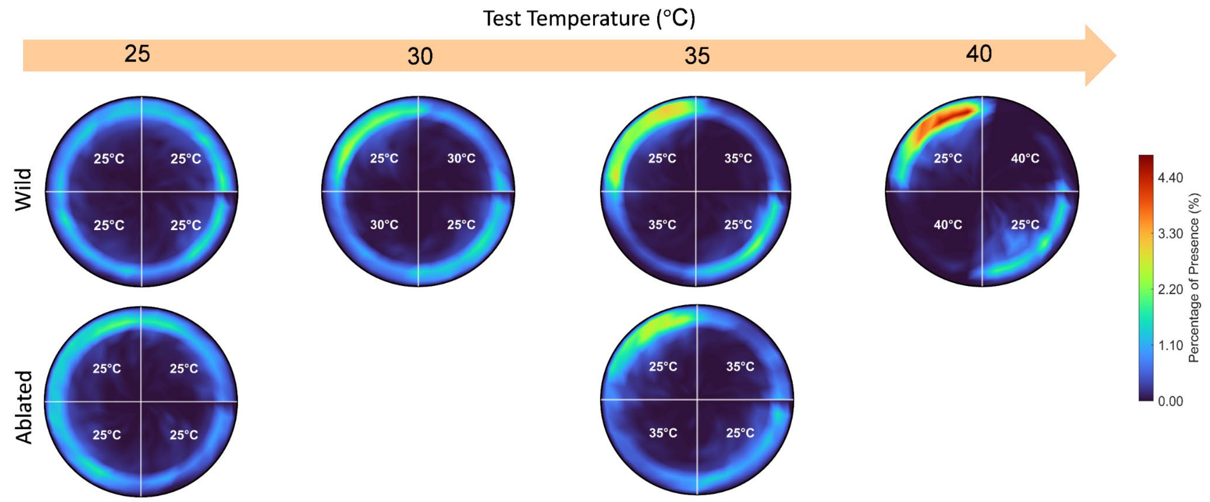


**Figure S 3. Temperature Preference of Wild-Type and Antennae-Ablated Groups in Two-Choice Assays for Quadrants I and III.** Consistent with observations from assays with test temperatures in quadrants II and IV, both wild-type and antennae-ablated groups showed a marked increase in heat avoidance behavior as temperature increased. These findings indicate that thermal preference is the dominant factor influencing navigational decisions, surpassing the effects of other experimental variables such as the spatial configuration of base and test zones or incidental fly-to-fly interactions.

### Braitenberg Model Parameters

**Table S 1. Parameters Used in the Braitenberg Vehicle Model Simulations.** This table summarizes the parameter values applied in the Braitenberg Vehicle Model for different vehicle starting positions. The model assumes that when the vehicle moves from the base region into the boundary region, inhibitory action prevents entry into the test region by applying a positive ipsilateral weight ($w_I$, excitatory) and a negative contralateral weight ($w_C$, inhibitory). These weights induce U-turns at the boundary, guiding the vehicle back to its original base quadrant. If the boundary is near the opposite base quadrant, the vehicle has an increased likelihood of entering that quadrant. Conversely, when moving from the test region to the boundary region, positive weights encourage quick entry into the base region. Sensor nonlinearity parameters (a=0.5 $\frac{1}{°C}$, b=3.9) remain constant across all regions. The initial velocity ($v_0$) is based on experimental data.

| Initial Region of the Vehicle | $w_I(\frac{mm}{s})$ | $w_C(\frac{mm}{s})$ | $v_0(\frac{mm}{s})$ |
|---|---|---|---|
| Base Region | 3 | 18 | 4.14 |
| Test Region | 20 | 30 | 6.44 |
| Boundary Region – From Same Base Quadrant (U-turn) | 18 | -30 | 4.63 |
| Boundary Region – From Opposite Base Quadrant | 1 | 20 | |
| Boundary Region – From Test Quadrant | 1 | 14 | |
